# Pools of independently cycling inositol phosphates revealed by pulse labeling with ^18^O-water

**DOI:** 10.1101/2024.05.03.592351

**Authors:** G. Kim, G. Liu, D. Qiu, N. Gopaldass, G. De Leo, J. Hermes, J. Timmer, A. Saiardi, A. Mayer, H.J. Jessen

## Abstract

Inositol phosphates control many central processes in eukaryotic cells, including nutrient availability, growth, and motility. Kinetic resolution of a key modulator of their signaling functions, the turnover of the phosphate groups on the inositol ring, has been hampered by slow uptake, high dilution, and constraining growth conditions in radioactive pulse-labeling approaches. Here, we demonstrate rapid (seconds to minutes), non-radioactive labeling of inositol polyphosphates through ^18^O-water in yeast, amoeba and human cells, which can be applied in any media. In combination with capillary electrophoresis and mass spectrometry, ^18^O-water labeling simultaneously dissects the *in vivo* phosphate group dynamics of a broad spectrum of even rare inositol phosphates. The improved temporal resolution allowed us to discover vigorous phosphate group exchanges in some inositol poly- and pyrophosphates, whereas others remain remarkably inert. Our observations support a model in which the biosynthetic pathway of inositol poly- and pyrophosphates is organized in distinct, kinetically separated pools. While transfer of compounds between those pools is slow, each pool undergoes rapid internal phosphate cycling. This might enable the pools to perform distinct signaling functions while being metabolically connected.

## Introduction

Water soluble cytosolic inositol phosphates (InsPs) can be synthesized by rearrangement of glucose-6-phosphate to inositol-3-phosphate or they emanate from the hydrolysis of inositol lipids (Figure 1). Inositol-containing metabolites have key signaling roles in eukaryotic cells. Phosphatidylinositol lipids (PtdInsPs or simply PIPs) determine the identity of organelles and organize vesicular traffic between them (Posor *et al*, 2022). The biologically occurring species with phosphate groups in different positions of the inositol headgroup are important to cell polarity, movement and mechanotransduction, the release of growth factors, neurotransmitters, and hormones (Falkenburger *et al*, 2010). The cytosolic or water-soluble InsPs have equally diverse and important signaling functions, including DNA stability, RNA transport, transcriptional control, MAP kinase signaling, energy and phosphate homeostasis, and many others (Monserrate & York, 2010; Shears *et al*, 2012; Maffucci & Falasca, 2020). They phosphate homeostasis to ATP levels across species, in part by signaling via SPX domains (Li *et al*, 2020; Moritoh *et al*, 2021; Wild *et al*, 2016; Szijgyarto *et al*, 2011; Gu *et al*, 2021; Austin & Mayer, 2020).

**Fig. 1:**
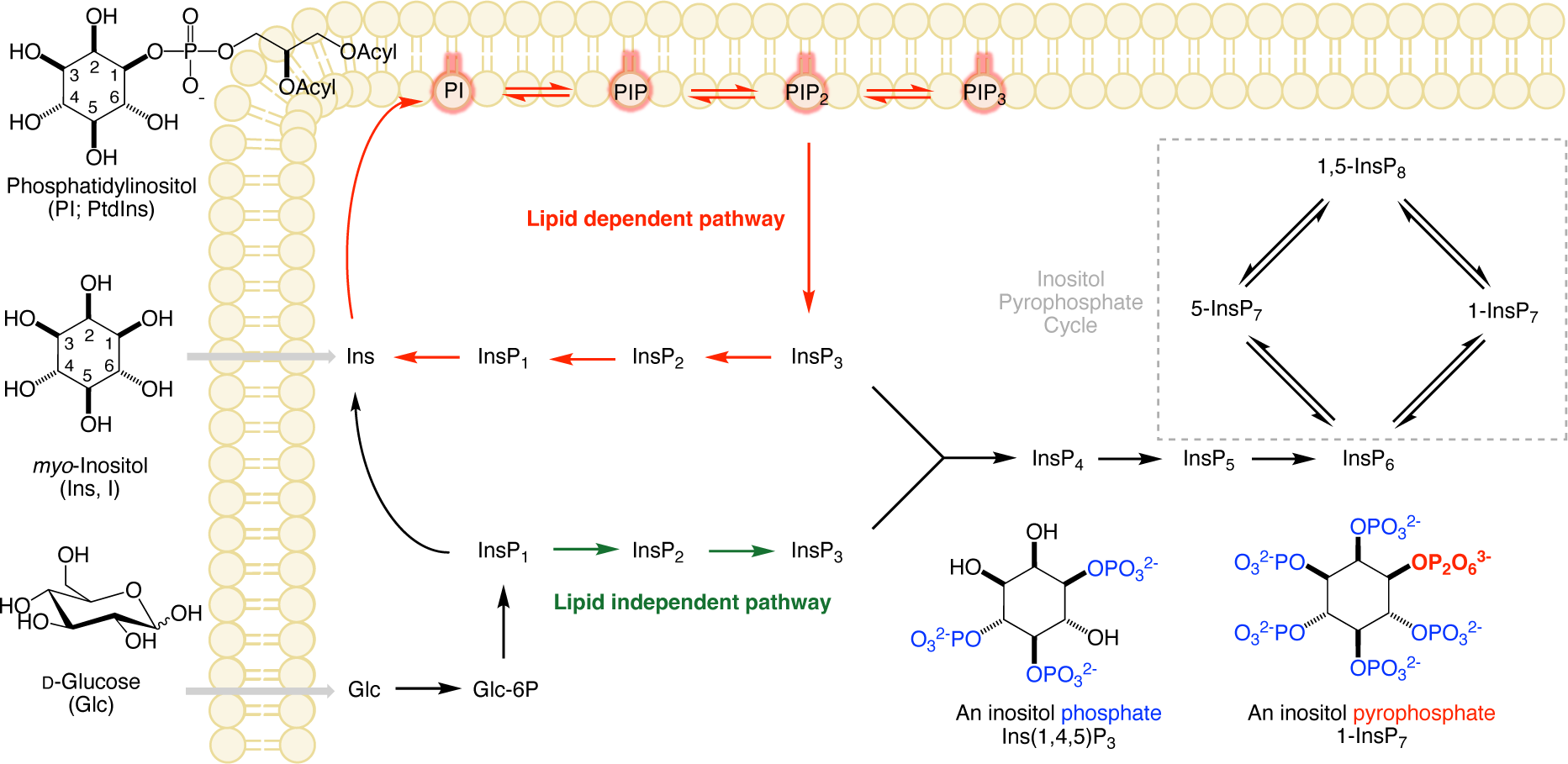
Pathways for synthesis of lipidic and generic soluble inositol phosphates. Inositol can either be taken up by cells or synthesized from glucose after uptake and phosphorylation. Cytosolic inositol phosphates are generated from PtdInsP_2_ through phospholipase C, generating diacylglycerol and InsP_3_ (Berridge & Irvine, 1984). In mammalian cells, an additional lipid-independent pathway can generate InsP_3_ directly from glucose (Desfougères *et al*, 2019). Forward phosphorylation by kinases up to InsP_8_ is possible while phosphatases remove the phosphates groups, generating lower phosphorylated forms. The balance between kinases and phosphatases will likely occur at different rates controlled by metabolic and/or signaling inputs. Therefore, we can envisage an InsPs network constantly turning over, harmonizing the steady-state concentrations of the different metabolites.

In many of the above-mentioned processes, InsPs and PtdInsPs are interconverted, leading to alterations of the number and/or the positioning of their phosphate groups. Measuring the steady state levels of InsPs and PP-InsPs has usually been achieved by strong anion exchange (SAX) HPLC after metabolic labeling with ^3^H or ^14^C-inositol, but this is a lengthy process that requires 24 hours in yeast to several days in mammalian cells under constraining growth conditions (Wilson & Saiardi, 2017). More recently, capillary electrophoresis mass spectrometry (CE-MS) has been introduced to monitor absolute InsP and PP-InsP levels (Eisenbeis *et al*, 2023; Liu *et al*, 2023; Qiu *et al*, 2020), and stable isotope ^13^C labeling with glucose and inositol has revealed alternative pathways of soluble InsP synthesis from membrane lipids or glucose (Figure. 1) (Desfougères *et al*, 2019; Trung *et al*, 2022). Although these approaches can be used to separate and quantify products of the steady state levels of InsPs and PP-InsPs, the fast dynamics of the interconversion of the phosphate groups has remained difficult to capture. This is in large part due to inherent limitations and complexity of the pulse-labeling approaches with radiolabeled tracers, which have so far been necessary for such analyses.

Inositol phosphates can be labeled through feeding of the cells with radiolabeled inositol, or by adding isotope-labeled phosphate (Wilson & Saiardi, 2017). Inositol uptake is slow, and labeling is usually performed to isotopic equilibrium, involving culture in inositol-depleted media over several generations. Moreover, inositol can be synthesized by cells from glucose, which leads to unlabeled inositol that cannot be traced (Qiu *et al*, 2020; Desfougères *et al*, 2019). Phosphate labeling is also comparatively slow because it relies on uptake through plasma membrane transporters. It is subjected to isotopic dilution both in the medium, where phosphate is an essential macronutrient, and in the cell, which has cytosolic phosphate concentrations of up to 20 mM (Auesukaree *et al*, 2004) and storage pools for phosphate. Furthermore, P_i_ utilization, and thereby the ability to label the internal P_i_ pool through P_i_ uptake, is strongly influenced by biosynthesis and cell growth, which change as a function of the culture conditions.

While steady state levels of metabolites themselves are an important information, the dynamic turnover of metabolites in this steady state, the metabolic flux, reflects the activity of a metabolic pathway (Nemutlu *et al*, 2012). Flux provides additional information about the metabolic state and can constitute a signal for the cell (Kotte *et al*, 2010; Kochanowski *et al*, 2013; Nemutlu *et al*, 2015). It can help us understand how a steady state concentration is maintained and how quickly a system can adjust to perturbations (Monge *et al*, 2019; Dawis *et al*, 1989; Desfougères *et al*, 2019). For example, the use of NaF as metabolic trap to block metabolic fluxes depending on phosphatases revealed dynamic phosphorylation of the abundant InsP_6_ (50-100 µM) into scarce PP-InsPs (0.1 - 2 µM), with a turnover of ca. 50% of the whole InsP_6_ pool within one hour (Menniti *et al*, 1993).

Measuring dynamic turnover of phosphorylated metabolites should ideally rely on direct labeling of the phosphate groups. This can be achieved by isotopes. For phosphorous, only radioactive nuclides are available. Furthermore, phosphorous can be absorbed by cells only in the form of organic or inorganic phosphate (P_i_). The uptake process can be quite slow, rendering it difficult to monitor rapid processes. Another option of labeling the phosphate group is by stable isotope labeling using ^18^O-enriched water (Dawis *et al*, 1989; Wang *et al*, 2021; Barneda *et al*, 2022). Water enters cells within seconds. It is incorporated into P_i_ by hydrolytic enzymatic reactions. A purely chemical exchange of P_i_ is inefficient (Dawis *et al*, 1989; Hackney, 1980). Hydrolysis occurs vigorously on cellular nucleotides, as exemplified by ATP, for which the entire cellular pool can turn over in seconds (Dawis *et al*, 1989; Versaw & Metzenberg, 1996). This rapid cycling entrains a similarly rapid introduction of ^18^O into the hydrolytic product, P_i_.

The incorporation of ^18^O into ATP (9 oxygens are exchangeable via different enzymatic mechanisms) has been studied extensively using direct ^31^P NMR approaches and LC-MS, as well as by an indirect GC-MS approach (Juranić *et al*, 2011; Nemutlu *et al*, 2012). Due to the inherently low sensitivity of NMR, this approach will not be effective to monitor scarce signaling molecules with high turnover rates. Of particular interest for phosphotransfer reactions is the appearance of the isotope label in the γ-phosphate of ATP, as this is the most readily transferable phosphate group. In rat hearts, for example, the γ-phosphate in every ATP is turned over 34 times per minute, the β-phosphate 12 times and the α-phosphate only once a minute (Nemutlu *et al*, 2012). ATP-synthase can incorporate ^18^O from ^18^O-water even multiple times into ATP before releasing its product into the cell (Walker, 1998; Dawis *et al*, 1989). Earlier experiments with ^18^O-water have also underpinned the existence of different compartmentalized ATP pools (metabolic and nonmetabolic), which vary with cell types and physiological states (Dawis *et al*, 1989). While ^18^O-labeling was exploited for a limited number of analyses of ATP turnover, and labeled synthetic ^18^O ATP has been used in phosphoproteomics (Müller *et al*, 2016; Li *et al*, 2014), the potential of an extension of this concept to other areas of phosphorylated metabolites received only limited attention. Recently, a study has shown incorporation of ^18^O labels into the PtdInsPs phosphate diester after incubation of mammalian cells with ^18^O-water (Barneda *et al*, 2022). Furthermore, the advent of new chemical synthesis approaches made ^18^O labeled reference compounds available, enabling ionization and fragmentation studies to improve assignments and quantifications of inositol phosphates (Haas *et al*, 2021; Hofer *et al*, 2015; Qiu *et al*, 2022).

Here, we use pulse-labeling to explore the turnover of inositol polyphosphates of yeast, amoeba and mammalian cells. Since these are in large part low-abundance metabolites (nanomolar to low micromolar concentration), we rely on the sensitivity and separation power of capillary electrophoresis coupled to both QQQ and qTOF mass spectrometry for their analysis. Our *in vivo* approach is based on the rapid permeation of ^18^O water into cells, and on the extremely rapid turnover of the cellular P_i_ through nucleotide hydrolysis (Versaw & Metzenberg, 1996; Heijnen, 2010). Our results provide an entry point into coupled InsP and PP-InsP fluxomics, revealing unexpectedly high turnover rates of some metabolites, metabolic lethargy of others, and the existence of several independently cycling InsP pools.

## Results

### Rapid ^18^O labeling of ATP in yeast

To label yeast cells, cultures that were logarithmically growing in SC (synthetic complete) medium were transferred to the same SC medium made with 50% ^18^O-water. 50% ^18^O-water suffices to label the entire pool of P_i_ almost quantitatively because, in isotopic equilibrium, the probability of P_i_ remaining with four unlabeled oxygen atoms is equal to 1/16 (6%). In this medium, P_i_ is the sole source of phosphate. At different time points, aliquots of the culture were extracted with perchloric acid (Wilson *et al*, 2015; Wilson & Saiardi, 2018) and the extracts were analyzed by CE-MS. The concentration of ATP was determined by comparison to synthetic ^13^C-ATP, which had been used as internal standard. The number of labels was assigned by combination of internal standard and high resolution qTOF mass spectrometry (Figure 2). Already at the first time point taken, after 1 min, ATP with one to four ^18^O atoms was detected (Figure 2). Labeling proceeded rapidly, reaching up to seven ^18^O labels after 60 min. The ^18^O labels also rapidly penetrated the InsPs, which could be analyzed from the same extracts as used for nucleotide analytics. The gradual incorporation of multiple labels poses a problem for rare analytes because the overall abundance of ions is now distributed over all isotopologues. This renders the MS analysis of rare analytes challenging, such as InsP_5_, InsP_7_ and InsP_8,_ and complicates detection of analytes with a large variable number of ^18^O atoms. To try to quantify all these rare analytes, CE-MS with a triple quadrupole system in multiple reaction monitoring (MRM) mode was employed, which has higher sensitivity as compared to a qTOF system.

**Fig. 2:**
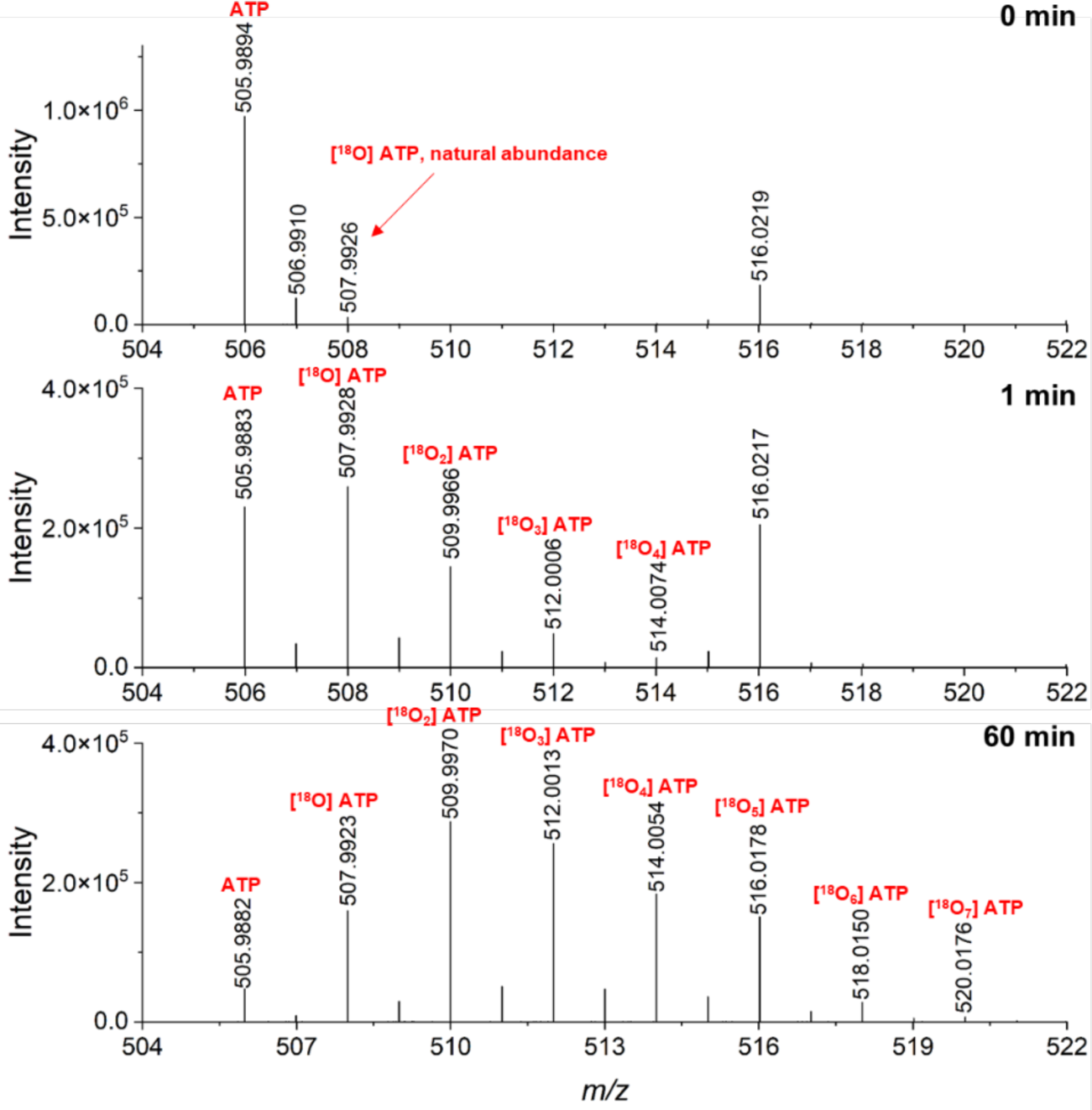
qTOF MS analysis of ATP. The analysis reveals the kinetics of ^18^O incorporation into ATP at different exchangeable positions. Wild-type yeast cells were grown in SC medium at 20°C to slow down the incorporation. At the 0 min timepoint, the medium was changed to SC medium prepared with 50% ^18^O-water. Samples were harvested at different timepoints (1 min and 60 min), extracted with perchloric acid and TiO_2_ and analyzed by qTOF mass spectrometry. Theoretical [M-H]^-^ for ATP, [^18^O] ATP, [^18^O_2_] ATP, [^18^O_3_] ATP, [^18^O_4_] ATP, [^18^O_5_] ATP, [^18^O_6_] ATP and [^18^O_7_] ATP are 505.9885, 507.9927, 509.9970, 512.0012, 514.0055, 516.0097, 518.0139, 520.0182, respectively.

The number of labels on the γ-phosphate (i.e. the group that will be transferred to InsPs) was assigned by MS/MS under conditions where the γ-phosphate was cleaved off. For example, one of the major fragmented ions found was 408.0117 (corresponding to [M-H-H_3_PO_4_]^-^) for unlabeled ATP (Supplementary Figure S1). An unexpected oxygen exchange (scrambling) on ATP was detected through analysis of a synthetic control compound - γ-^18^O_2_ labeled ATP (Hofer *et al*, 2015) - which carries two ^18^O as non-bridging oxygens of the γ-phosphate (Supplementary Figure S2). Its fragmentation showed partial scrambling of ^18^O in the gas phase under these conditions, i.e. we observed mono-labeled γ-phosphate and mono-labeled ADP. Therefore, our approach bears the risk to underestimate the true amount of ^18^O labeling in the γ-phosphate. It is mechanistically conceivable that such an exchange of oxygens might occur through a reversible ping-pong gas-phase reaction involving metaphosphates. This interesting facet will be studied in more detail in a future study. Scrambling was also observed when synthetic [^18^O_2_] 5-InsP_7_ was employed (Haas *et al*, 2021), which was only labeled on the non-bridging oxygens in the β-phosphate (Supplementary Figure S3). To accurately count the numbers of ^18^O labels in the analytes and avoid an underestimation of the extent of labeling on ATP and InsPs, scrambling ion products had to be considered using the MRM transitions shown in Supplementary Table S3, Table S4, Table S5, Table S6, Table S7 and Table S8.

### Rapid pulse-labeling resolves separate, cycling pools of soluble inositol polyphosphates in yeast

Using our previously described CE-QQQ method (Qiu *et al*, 2020), we monitored the time course of ^18^O incorporation from ^18^O-water into ATP in general, but particularly the γ-phosphate of ATP. Already after 1 min more than 68% of the ATP pool was labeled (Figure 3A, for representative example of extracted ion electropherograms see Supplementary Figure S4). When we focused the analysis only on the γ-phosphate of ATP, the kinetics were similarly rapid, with 43% carrying ^18^O within 1 min. The labeling tends to be even faster and is shifted to larger numbers of total ^18^O incorporation when 100% ^18^O-water was used in the culture medium (Figure 3B), which was avoided for cost reasons. Under these conditions, 90% of the γ-phosphate of ATP are stably labeled within 5 minutes (Figure 3B).

**Fig. 3:**
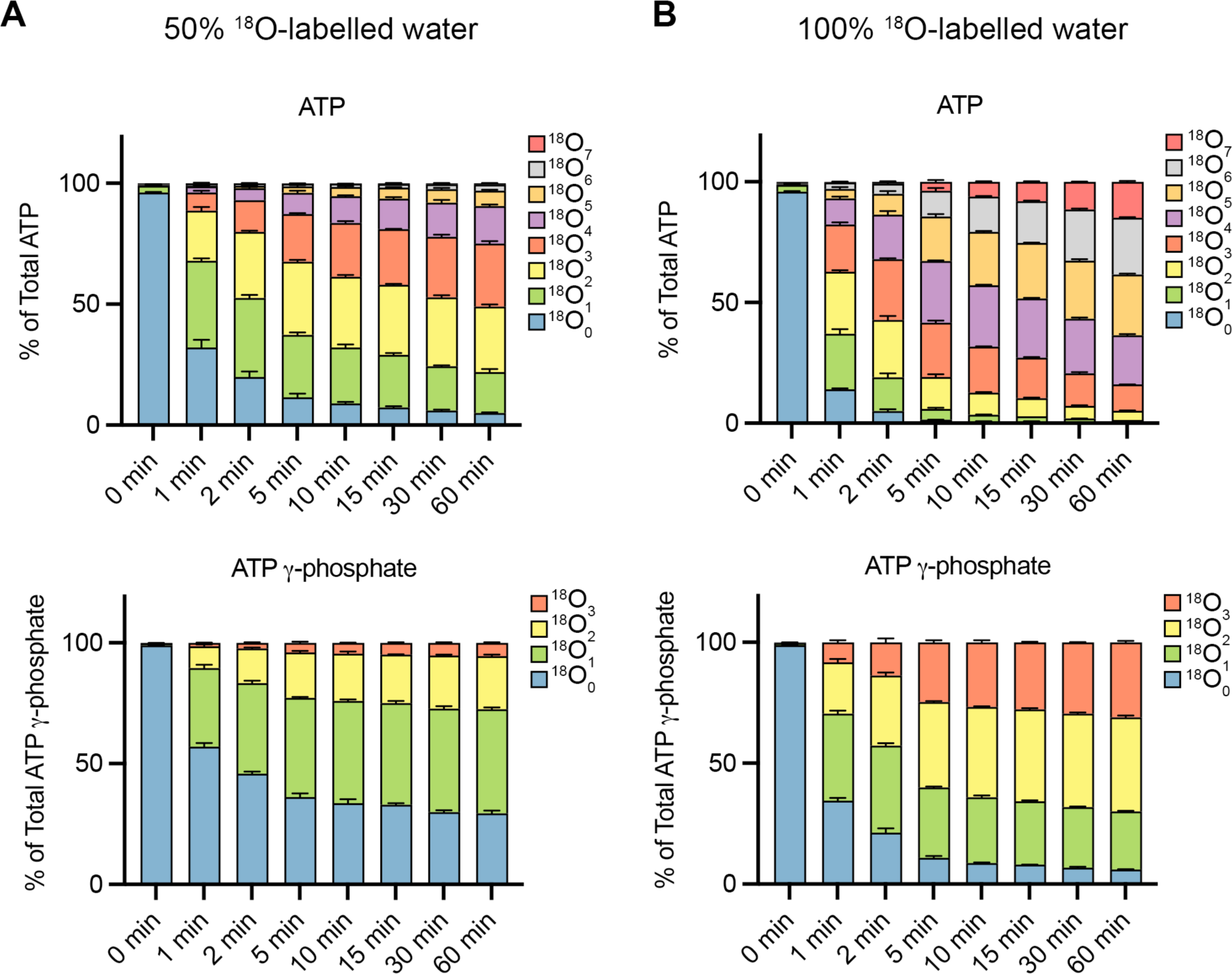
Kinetics of ^18^O entry into ATP of yeast under steady state conditions. Time-dependent formation of ATP isotopologues in yeast with different numbers of ^18^O atoms studied by CE-QQQ. Wild type yeast cells were grown logarithmically in SC medium at 20°C. At the 0 min time point, the medium was changed to SC medium prepared with **A** 50% or **B** 100% of ^18^O-labelled water (99% enrichment). After further incubation at 20°C for the indicated periods of time, cells were extracted with perchloric acid and TiO_2_. The graphs show the fractions of ATP that carried the indicated numbers of ^18^O at any position (top panels) or in the γ-phosphate (bottom panels). The means of 3 samples with standard deviation are shown.

The ^18^O label rapidly appeared in the InsPs and PP-InsPs. Our measurements indicated that InsP_7-8_ already contained ^18^O at the 0 min time point above the expected natural abundance of ^18^O (isotope frequency is 0.2% for each oxygen). As inositol polyphosphates are very oxygen-rich molecules (e.g. 27 oxygens for InsP_7_), this leads to an expected fraction of ca. 5 % of these compounds carrying at least one ^18^O. Our analytical protocol overestimates this expected value due to isotopic peaks of unlabeled InsPs using the wide mass resolution (full-width at half maximum of 1.2 Da, ± 0.6 Da mass error) of the quadrupole that we had to choose in these experiments. Wide mass resolution can separate major isotopic clusters, such as carbon-based compound dominated by ^12^C and ^13^C, referred to as M and M+1. In the case of InsP_7_ and InsP_8_, doubly charged precursor ions and product ions were detected, resulting in a skewing of the expected value by ca. 3-fold at the 0 min time point. This skewing could be corrected by performing unit mass resolution, which is specified to have ± 0.35 Da mass error (Supplementary Figure S5). However, unit mass resolution was not sufficiently sensitive to quantify rare analytes, such as ^18^O labeled InsP_7_ and InsP_8_. To estimate whether the described skewing effect changed its magnitude during the time course, mixtures of synthetic ^18^O_2_ 5-InsP_7_ and unlabeled 5-InsP_7_ with different compositions were prepared and analyzed by CE-QQQ using the wide mass resolution. Raising the concentration of the ^18^O_2_ 5-InsP_7_ leads to a corresponding decrease of the skewing of ^18^O 5-InsP_7_ (Supplementary Figure S6). The effect is largest at the zero-time point (ca. 10%) in our experiments and declines during increasing incorporation of ^18^O labels, which made a simple subtraction to the expected zero-time point value problematic. A two-fold change in the concentration of ^18^O_2_ 5-InsP_7_ was accompanied by a skewing decrease of ca. 2%. Therefore, we present overestimated ^18^O labeling values of InsP_7_ and InsP_8_ in this study.

When the cells were transferred from unlabeled culture medium into medium of the same composition but made from ^18^O-water (50%), 32% of 5-InsP_7_ and 44% of 1,5-InsP_8_ acquired one or two ^18^O labels already within 1 min (Figure 4A and B, for representative example of extracted ion electropherograms see Supplementary Figure S7). For 1,5-InsP_8_, this led to 78% labeling of the pool within 15 min and remained almost constant thereafter, whereas labeled 5-InsP_7_ maintained at 66% after 60 min. This suggests the existence of at least two pools of 5-InsP_7_, one resting static and the other one turning over very vigorously. 100% ^18^O-water labeling results confirmed our hypothesis by showing complete labeling of the 1,5-InsP_8_ pool within 30 min while 5-InsP_7_ maintained a plateau at 60% (Supplementary Figure S8). In contrast to the inositol pyrophosphates, InsP_6_ was labeled only very slowly, showing ca. 7% of ^18^O labeling after 60 min (Figure 4C). This suggests that the InsP_6_ pool is relatively static regarding its synthesis from InsP_5_, whereas the inositol pyrophosphates undergo highly dynamic, permanent turnover of their phosphate groups attributable almost exlusively to exchange of the β-phosphates of the P-anhydrides. Remarkably, this high turnover occurs even under constant nutrient availability. It hence appears to be a constitutive feature of inositol pyrophosphate metabolism. Notice that the turnover of PP-InsPs must include turnover of InsP_6_, but since no labels appear rapidly in InsP_6_, this is due to synthesis and cleavage of the diphosphate groups.

**Fig. 4:**
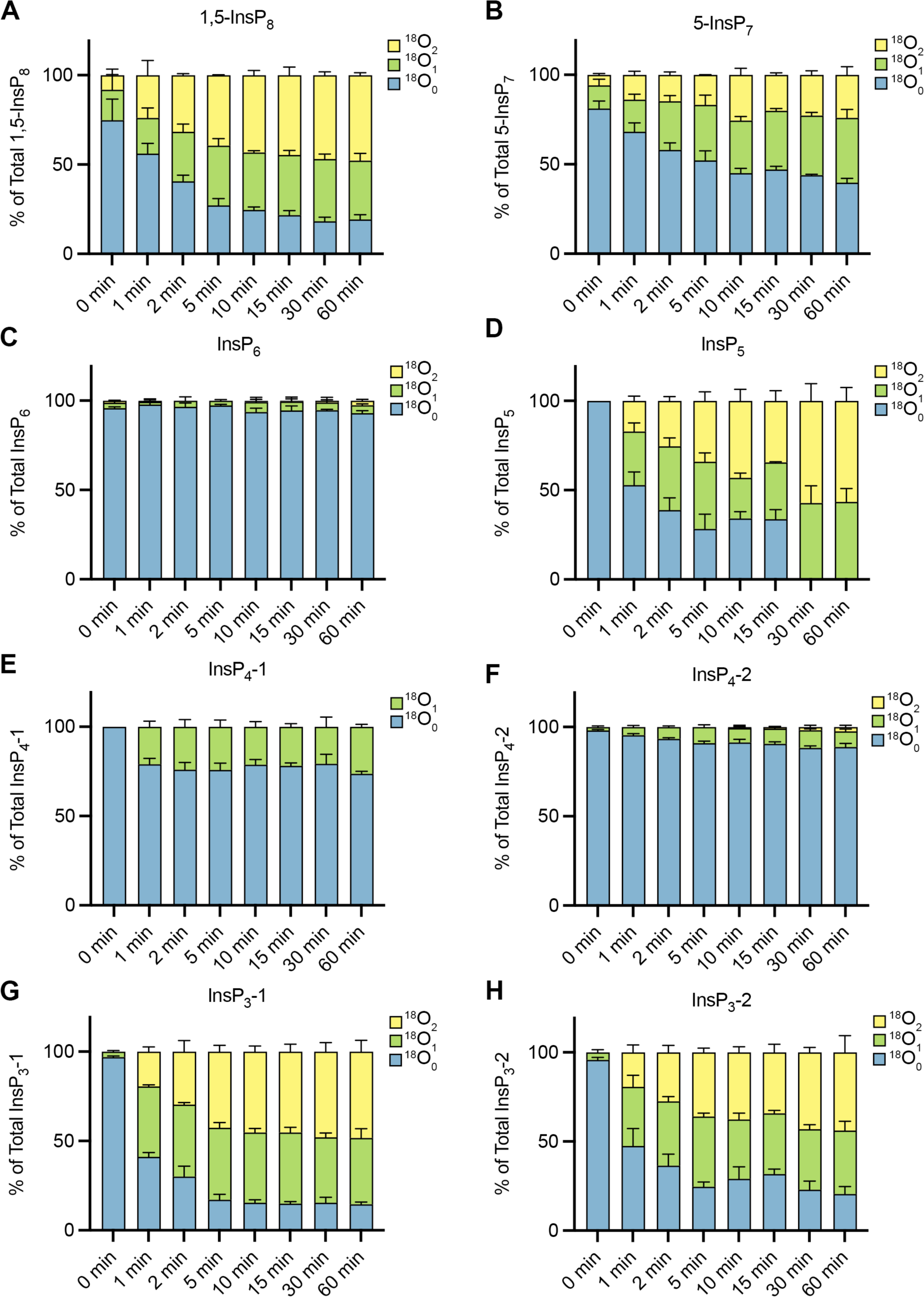
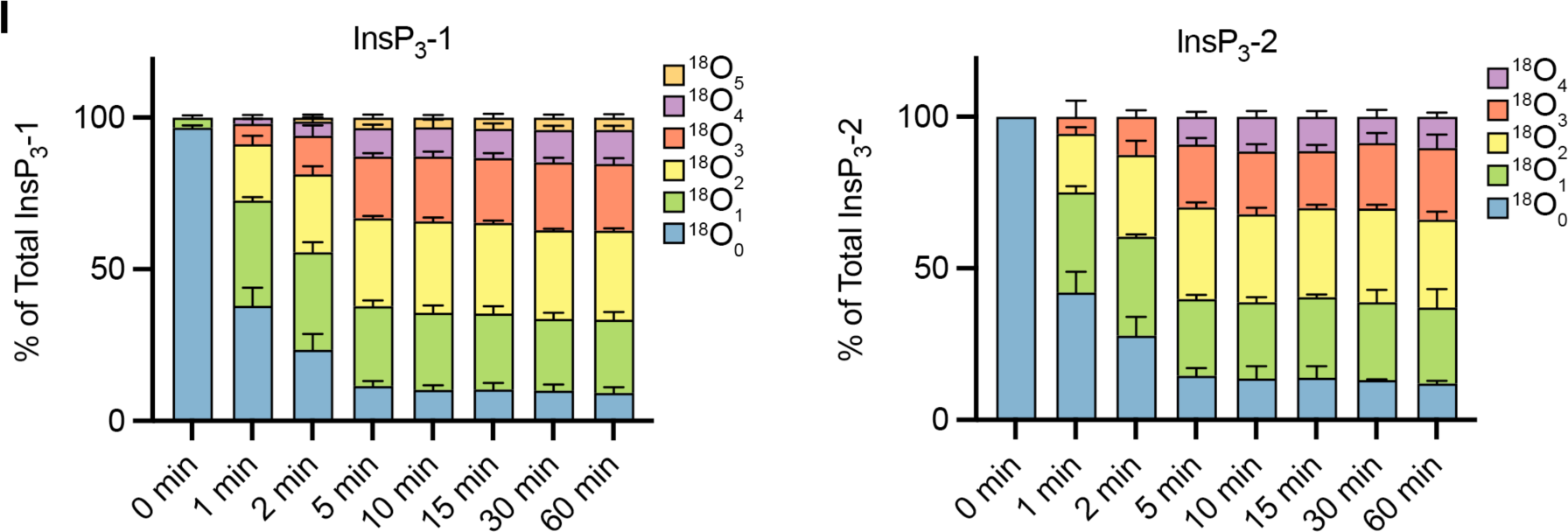
Kinetics of ^18^O entry into soluble InsPs of yeast under steady state conditions. Wild type yeast cells were grown logarithmically in SC medium at 20°C. The medium was changed to SC medium prepared with 50% of ^18^O-labelled water. After further incubation at 20°C for the indicated periods of time, cells were extracted with perchloric acid and TiO_2_ and analyzed by CE-QQQ. Note that the levels of 1-InsP_7_ in these samples were too low to be reliably detected. The means of 3 samples with standard deviation is shown, representing **A**. 1,5-InsP_8_, **B**. 5-InsP_5_, **C**. InsP_6_, **D**. 2-OH-InsP_5_, **E**. InsP_4_-1, **F**. InsP_4_-2, G. InsP_3_-1, **H**. InsP_3_-2: Ins(1,4,5)P_3_, **I**. Higher numbers of ^18^O incorporation on InsP_3_ were observed by CE-qTOF of samples as in G and H.

The analysis of lower phosphorylated InsPs using the same experimental approach revealed further interesting behavior. InsP_3_, which is generated from PI(4,5)P_2_ through phospholipase C (Figure 1), incorporated ^18^O into ca. 50% of the pool within one minute (Figures 4G and 4H). The InsP_3_ pool could be dissected into two different species with similar dynamics. One of these species has the same electrophoretic migration time as Ins(1,4,5)P_3_ (assigned as InsP_3_-2 through comparison with a ^13^C-labeled internal reference, Supplementary Figure S9), which thus represents the direct product of PIns(4,5)P_2_ hydrolysis. The other InsP_3_ (assigned as InsP_3_-1) does not comigrate with any reference InsP_3_ isomers that are available in our laboratory. We solely do not have an Ins(2,4,6)P_3_ reference, and thus InsP_3_-1 most likely represents Ins(2,4,6)P_3_. Interestingly, the incorporation of up to five ^18^O labels into InsP_3_ were detected by CE-qTOF, suggesting that several phosphate groups must undergo exchange. This is likely a result of the highly dynamic lipid dependent turnover of Ins(1,4,5)P_3_ but cannot explain the phenomenon for Ins(2,4,6)P_3_ which does not derive from this lipid (Figure 4I, and Supplementary Figure S10).

InsP_4_ showed comparably limited turnover (Figure 4E and 4F). Also InsP_4_ could be dissected into two different InsP_4_ species. One of them was static after 1 min and incorporated ^18^O to ca. 20%, the other one only showed very sluggish ^18^O incorporation. The different pools of InsP_4_ very likely represent positional isomers with regards to the phosphate groups and not diastereomeric inositol core structures, such as *scyllo* inositol. We have not assigned the identities of these species further within this study.

### Kinetic compartmentalization of the InsP pathway in human cells

Next, we tested the suitability of ^18^O-water to label inositol phosphates in mammalian HCT-116 cells. These experiments employ larger volumes of media and were hence performed with 50% ^18^O-water to reduce cost. Cells were pulse-labeled by transferring them into medium made with 50% ^18^O-water but of otherwise identical composition. In contrast to the yeast experiments, we incubated at 37°C in a 5% CO_2_ atmosphere. At different time-points of pulse-labeling, aliquots were extracted by perchloric acid and analyzed by CE-MS as described above (Figure 5).

**Fig. 5:**
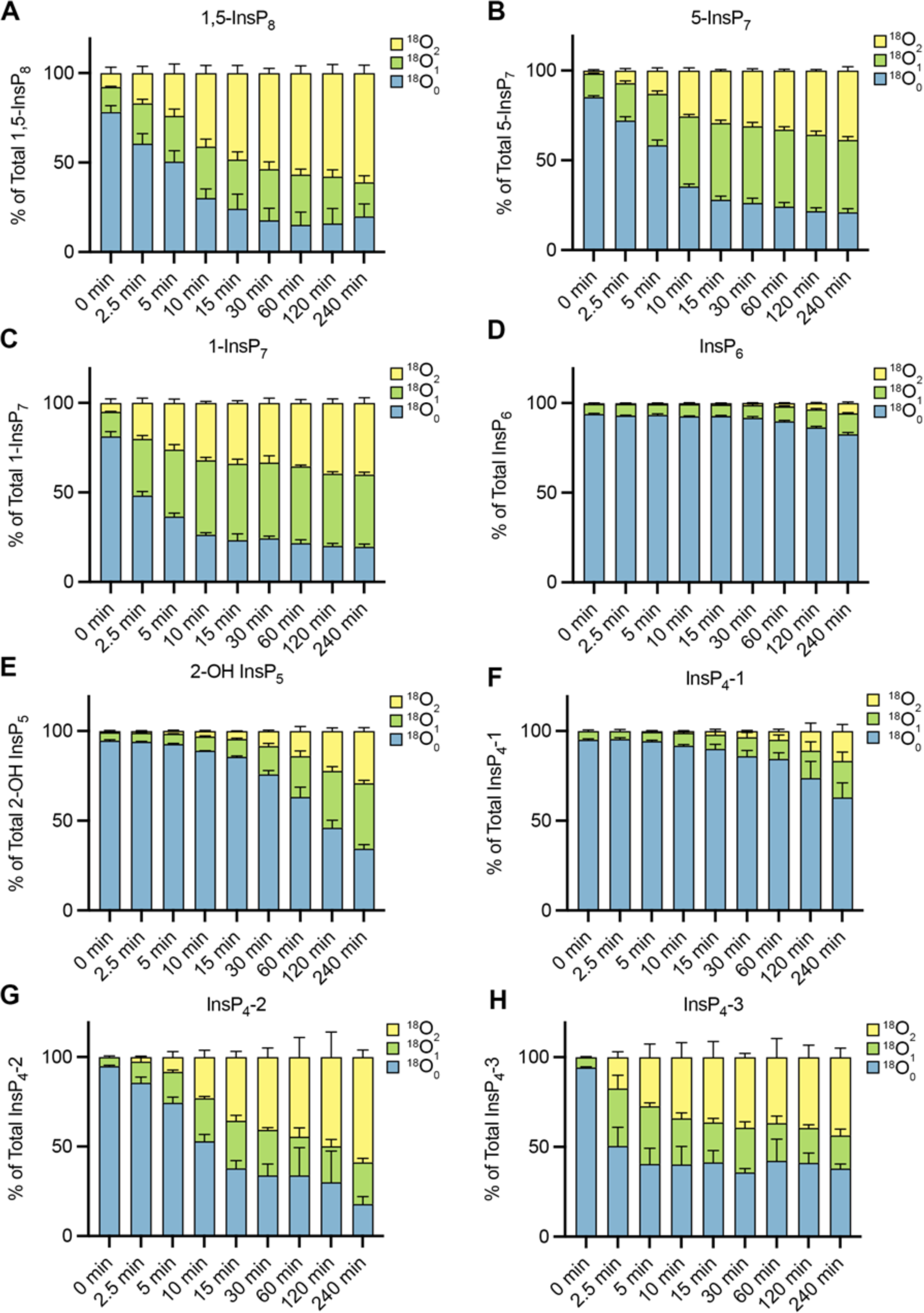
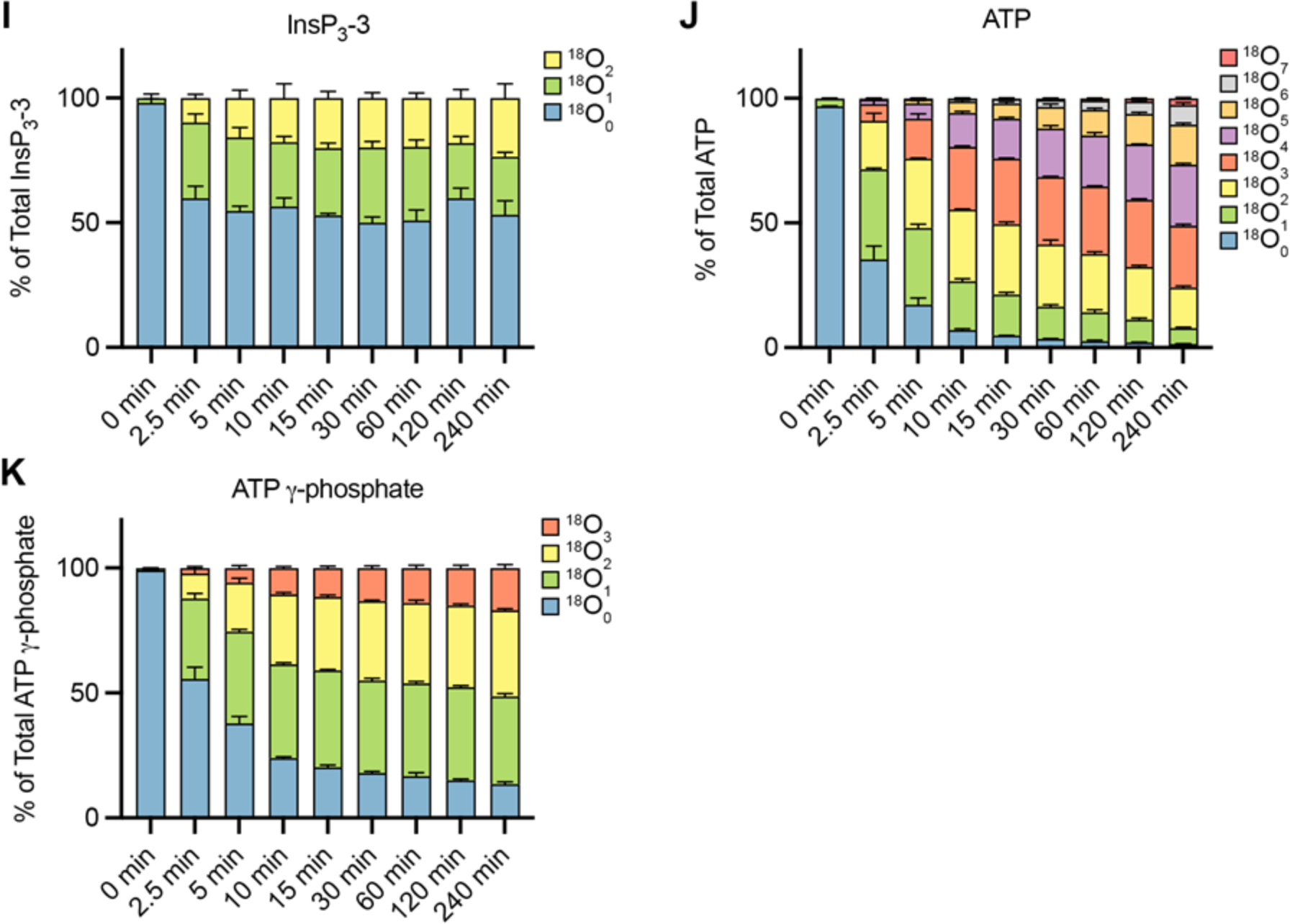
Kinetics of ^18^O entry into ATP and cytosolic InsPs from human cells. Kinetics of ^18^O entry into 1,5-InsP_8_ (**A**), 5-InsP7 (**B**), 1-InsP_7_ (**C**), InsP_6_ (**D**), 2-OH InsP_5_ (**E**), InsP_4_-1 (**F**), InsP_4_-2 (**G**), InsP_4_-3 (**H**), InsP_3_-3 : Ins(1,4,5)P_3_ (**I**), ATP (**J**) and the γ-phosphate of ATP (**K**) were monitored in mammalian cells. HCT116 cells were grown in DMEM medium, detached and collected by centrifugation. The cell pellet was resuspended in DMEM medium prepared with 50% of ^18^O-labeled water. After the indicated periods of further incubation at 37°C, aliquots of the cells were extracted with perchloric acid. CE-MS analyses of the extracts were performed. The means of three replicates are shown with standard deviations.

^18^O rapidly entered ATP and the inositol pyrophosphates (5-InsP_7_; 1-InsP_7_; 1,5-InsP_8_), reaching an equilibrium after 10-20 min. Between 20 and 25% of these PP-InsPs remained unlabeled. InsP_6_ labeling proceeded slowly also in mammalian cells, reaching only 1/6^th^ of the pool after 4 hours. This incorporation probably reflects new synthesis of InsP_6_ through cell growth because the generation time of the cells under these steady media conditions is around 24h. During the 4h of labeling time, biomass would thus increase by roughly 1/6^th^, accounting for the increase in labeled InsP_6_, which is of the same order. Interestingly, 2-OH InsP_5_, which is of similar abundance as InsP_6_ in some mammalian cells (Qiu *et al*, 2022), showed significantly faster labeling than InsP_6_, suggesting that the flux from InsP_5_ to InsP_6_ by the inositol-pentakisphosphate 2-kinase (IPPK) is sluggish. The appearance of only two labels suggests that turnover takes place at a specific position and not at many different ones, as this would lead to higher numbers of ^18^O incorporations.

Each InsP_4_ isomer showed different ^18^O incorporation patterns. InsP_4_-1 and InsP_4_-2 incorporated ^18^O at constant rates but with quite different kinetics. InsP_4_-1 exhibited relatively slow incorporation of ^18^O, similar to InsP_6_, whereas 80% of InsP_4_-2 was labeled with ^18^O over 4h. In contrast, InsP_4_-3 incorporated ^18^O very rapidly, reaching a plateau within 2.5 min that remained constant for the rest of the 4h incubation period. Unlike the minimal turnover of InsP_4_ observed in yeast, significant turnover was apparent in HCT-116 cells.

We could resolve three peaks of InsP_3_. The first two, InsP_3_-1 and InsP_3_-2, had no detectable ^18^O labels. We assigned InsP_3_-1 as Ins(1,2,3)P_3_ and InsP_3_-2 as Ins(1,2,6)P_3_ and/or its enantiomer Ins(2,3,4)P_3_ (identified by comparison with ^13^C-labeled internal references, Supplementray Figure S9). InsP_3_-1 and InsP_3_-2 are apparently not significantly generated through kinases, which would lead to incorporation of ^18^O label. They might, however, originate from dephosphorylation of InsP_6_ into these metabolites (see discussion). Since generation of Ins(1,2,6)P_3_ through dephosphorylation was recently described using ^13^C NMR labeling (Trung *et al*, 2022), Ins(1,2,6)P_3_ likely represents the enantiomeric identity of InsP_3_-2.

The third peak Ins(1,4,5)P_3_ (assigned to InsP_3_-3 through comparison with ^13^C-labeled internal references, Supplementary Figure S9) incorporated ^18^O very rapidly, and similarly InsP_4_-3, attaining a plateau within 2.5 min that remained constant for the rest of the 4h incubation. As in yeast, up to five ^18^O labels of the Ins(1,4,5)P_3_ were detectable by CE-qTOF within 15 min, suggesting turnover of multiple phosphate groups (Supplementary Figure S11). Unfortunately, we could not record these multiple ^18^O labels accurately beyond 15 min because of matrix effects in these sample sets. Labeling did not reach the entire pool of Ins(1,4,5)P_3_. Thus, similarly as argued above for yeast, there might be sub-pools of this compound, some of which remain quite static, and others that turn over rapidly and are responsible for the initially rapid yet limited integration of ^18^O into the total pool of Ins(1,4,5)P_3_.

Together, these data suggest massively different dynamics of phosphate exchange on InsPs also in mammalian cells: Highly dynamic phosphate cycling of the P-anhydrides in the inositol pyrophosphate pools (5-InsP_7_; 1-InsP_7_; 1,5-InsP_8_) and of phosphate esters in InsP_3_, likely as a result of the lipid dependent turnover and cleavage by PLC, is separated by much more inert pools of InsP_4_ and InsP_6_. InsP_5_ seems to interconvert to InsP_4_ and back to InsP_5_ again more rapidly. Thus, the inositol pyrophosphate and InsP_3_ pools may cycle independently from each other, connected only through slow biosynthetic reactions, or separated by compartmentalization.

### ^18^O-water pulse labelling reveals a sluggish PP-InsPs turnover in amoeba

Finally, we performed similar labeling and extraction experiments with 50% ^18^O-water with the slime mold *Dictyostelium discoideum* (Figure 6). This organism has played an important role in the discovery of PP-InsPs, as their concentrations are comparably high reaching sub-millimolar level (Pisani *et al*, 2014). It produces a range of PP-InsP isomers different from those found e.g. in yeast and mammals (Qiu *et al*, 2020; Desfougères *et al*, 2022). In *D. discoideum*, half of the ATP was labeled in its γ-phosphate within 10 min of incubation, and around 75% of total ATP had incorporated ^18^O (Supplementary Figure S12). These proportions changed only little in the subsequent 20 min, suggesting the existence of two pools of ATP that turns over at different velocities.

**Fig. 6:**
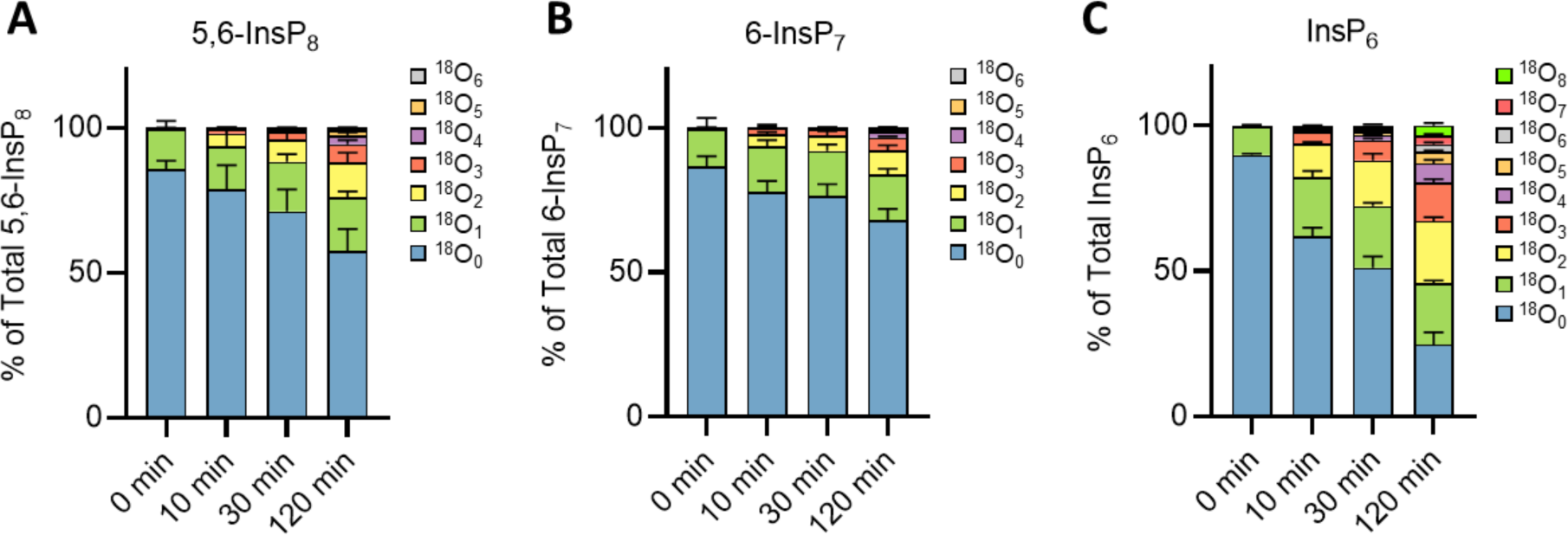
Kinetics of ^18^O entry into soluble InsPs from *D. discoideum*. Kinetics of ^18^O entry into 5,6-InsP_8_ **A**, 6-InsP_7_ **B**, InsP_6_ **C** monitored in *Dictyostelium*. Cells were pre-cultured in SIH medium and transferred to SIH media made of 50% of ^18^O-labeled water with P_i_. After further incubation for the indicated periods of time, samples were harvested and extracted. Extracts were measured by CE-QQQ. The means of four replicates with standard deviations are shown.

Labeling of InsP_7_ and InsP_8_ was much slower than in yeast and human cells: only 32% of InsP_7_ and 44% InsP_8_ incorporated labels over the full 2h of incubation time. Considering the approximately 8h generation time of *Dictyostelium*, the rate of incorporation into InsP_7_ may actually reflect new biosynthesis due to growth of the culture. The labeling of InsP_8_ goes beyond that but is far from the vigorous turnover observed in yeast and mammalian cells. InsP_6_ labeling, on the other hand, proceeded much faster than that of the inositol pyrophosphates, and it was also more rapid and more extensive than the labeling of InsP_6_ in yeast and human cells. Up to eight ^18^O labels were detected throughout the experiment, necessitating rapid phosphate group exchange at multiple positions in InsP_6_. Thus, the dynamics of InsPs metabolism in *Dictyostelium* are significantly different than in budding yeast or mammalian cells. Unfortunately, the sensitivity needed to obtain a comprehensive analysis of the labelling of InsP_5_, InsP_4_ and InsP_3_ was affected by the abundant and broad EDTA signal present from the extraction buffer. For this reason, we cannot yet report lower InsPs in these samples.

## Discussion

The metabolism of InsPs and PP-InsPs, which are signaling molecules of low abundance, has mainly been studied using radioactive labeling of inositol or of phosphate (Wilson & Saiardi, 2017). For inositol poly- and pyrophosphates, the labeling approach with ^18^O-water that we present here offers several important advantages, particularly for analyzing turnover of the phosphate groups (Versaw & Metzenberg, 1996; Mitchell *et al*, 1980; Barneda *et al*, 2022). ^18^O is a stable, non-radioactive isotope, such that media and samples produced with it can be stored for extended periods of time. ^18^O-water can be obtained in 99.5% pure form, enabling labeling to very high specific isotope content. This facilitates detection of inositol phosphates by mass spectrometry, some of which exist is sub-micromolar concentrations in cells (Chabert *et al*, 2023; Harmel *et al*, 2019; Barker *et al*, 2004; Albert *et al*, 1997). ^18^O-water labeling can be performed in any medium and under any growth condition. This is a distinctive advantage over labeling through radioactive inositol. Cellular membranes are also quite permeable to water, which allows very rapid labeling (seconds, as compared to hours for inositol labeling) to high specific isotope content and apparent equilibrium. In combination with CE-MS detection, this facilitated the analysis of unexpectedly rapid turnover of phosphate groups around the inositol ring with high specificity. Turnover can be monitored simultaneously for a variety of InsPs in the same experiment. The possibility of monitoring the kinetics of phosphate group turnover on InsPs with improved temporal resolution, sensitivity, throughput, and cost renders it possible to analyze flux through the metabolic pathways interconnecting inositol phosphates. It also allows to label inositol phosphate lipids. We exploit this to derive metabolic network models for inositol polyphosphate metabolism and to explore the dynamics of inositol lipids, aspects that will be described in separate studies.

Our analysis relies on the very high turnover of ATP inside cells, which renews the entire ATP pool at the timescale of seconds to a few minutes, depending on the cell type. Labeled γ-phosphates of ATP can then be transferred to inositol phosphates by respective kinases, and they can be removed again by hydrolysis through phosphatases. The efficiency of this ^18^O-water labeling approach is illustrated by the rapid labeling kinetics that we could uncover for some compounds, such as the PP-InsPs and some InsP_3_ species. Other pools of inositol phosphates, such as InsP_6_ and InsP_4_, incorporated ^18^O very slowly, suggesting at first sight very low turnover.

It is worthwhile to consider the enzymatic aspect of this process. Kinases transfer phosphate from ATP through a variety of mechanisms (Kenyon *et al*, 2012). A common feature is that the γ-phosphate of ATP undergoes an attack, for example by an OH group of the substrate to be phosphorylated. When ADP leaves, the oxygen constituting the link to the phosphorylated substrate stems from the unlabeled substrate, not from the ^18^O labeled γ-phosphate group. If the substrate is dephosphorylated again, labeled water attacks the phosphate group and the substrate leaves, keeping its unlabeled oxygen (Lassila *et al*, 2011). Label is thus introduced by the kinase reaction and withdrawn by the opposing phosphatase reaction. We may then expect that the substrate is ^18^O labeled only as long as it maintains the labeled phosphate group. Therefore, InsP_6_ should not be considered as a truly static pool. The rapid labeling of 5-InsP_7_ through InsP_6_ kinase requires cycling of phosphate on position 5 of the inositol ring and engages InsP_6_ as a substrate. Thus, there should be rapid cycling between InsP_6_ and 5-InsP_7_ by phosphorylation and dephosphorylation, but it does not leave a corresponding ^18^O trace in InsP_6_. Along the same line, we expect the labeling of 2OH-InsP_5_ to stem from the phosphorylation of InsP_4_ through inositol polyphosphate multikinase (IPMK). That this accumulating label is propagated only very inefficiently from 2OH-InsP_5_ into InsP_6_ argues for a sluggish exchange between InsP_5_ and InsP_6_ under the conditions we investigated. For similar reasons, we expect the interconversion of InsP_3_ and InsP_4_ to be slow.

These very pronounced differences in turnover suggest that phosphate cycling occurs on discrete compounds along the inositol phosphate metabolic pathway. This separates the rapidly interconverting InsP_7_ and InsP_8_ from sluggish interconversions between InsP_3_ and InsP_4_, and between InsP_5_ and InsP_6_. It suggests that, even though InsP_4_ through InsP_8_ derive from InsP_3_ and enzymes can interconvert all these compounds (Figure 1) (Tsui & York, 2010; Dyson *et al*, 2012; Desfougères *et al*, 2019), they do not operate as a simple linear pathway. Instead, InsPs appear to be organized into a series of separated phosphate cycles (Figure 7). In this way, each cycle is free to mediate independent signaling processes. The metabolic reactions connecting the cycles may then serve mainly to replenish or diminish substrate levels in the cycle, for example when growth or changing nutrient conditions may necessitate an adaptation of the signaling properties. Such a scenario arises for example upon phosphate replenishment after starvation, when the InsP_7_ and InsP_8_ pools become significantly expanded (Chabert *et al*, 2023; Dong *et al*, 2019; Li *et al*, 2020; Riemer *et al*, 2021).

**Fig. 7:**
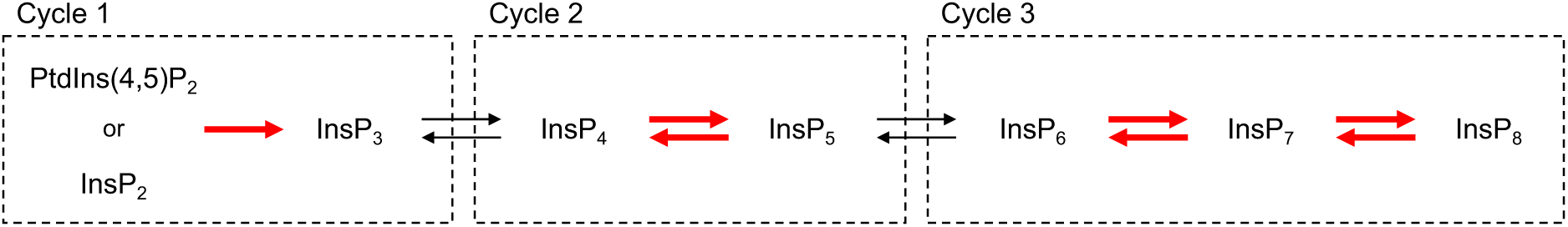
Three separate cycles of inositol phosphates. Inositol phosphates can be grouped into three distinct pools according to the turnover of their phosphate groups. Rapid interconversion of phosphate groups between metabolites is indicated by red arrows.

Turnover of PP-InsPs and InsP_3_ is very vigorous, even under steady growth conditions, i.e. under constant nutrient supply and growth. Such strong turnover implies permanent competing synthesis and degradation of the compounds. The simultaneous activity of two antagonizing reactions is often considered as "wasteful cycling". It may, however, be worth the price because it comes with significant benefits and additional potential for information transmission. Substrate cycling can sharpen the responses of a signaling system and increase its sensitivity. It can amplify signals (Goldbeter & Koshland, 1981, 1987), allow to filter or process information from noise in external and internal parameters, and create bistable switches (Samoilov *et al*, 2005; Miller & Beard, 2008). This makes for example the rapid cycling of 1,5-InsP_8_ interesting and potentially relevant because the yeast PHO pathway is controlled by 1,5-InsP_8_ (Chabert *et al*, 2023), exploits expression noise, and behaves like a bistable switch (Wykoff *et al*, 2007; Vardi *et al*, 2013, 2014; Raser & O’Shea, 2004).

Some inositol phosphates, such as 5-InsP_7_ in yeast or InsP_3_ in human cells, showed biphasic labeling kinetics, where a rapid initial turnover was followed by a long phase in which no or only negligible further ^18^O was incorporated. Since the cells had been growing under steady conditions, it is unlikely that this reflects a sudden change in the metabolism of these compounds. A more plausible explanation is sequestration of the compound into pools of greater or lesser accessibility to metabolizing enzymes. Such sequestration can be realized by stable binding of InsPs to proteins (Blind, 2020) or other molecules and/or by transfer into distinct subcellular compartments. For InsPs, several examples show that they can integrate into proteins in an apparently very stable fashion, such that the interaction persists even through purification and crystallization of the proteins. Examples include the capsid and Gag proteins of retroviruses (Obr *et al*, 2021; Renner *et al*, 2023) the RNA editing enzyme ADAR2 or casein kinase 2 (Macbeth *et al*, 2005; Lee *et al*, 2013). Moreover, InsPs might be sequestered into biomolecular condensate type structures such as nucleoli (Sahu *et al*, 2023).

Our profiling of InsP_3_ in human cells also revealed Ins(1,2,3)P_3_ and Ins(1,2,6)P_3_ and/or its enantiomer. Their undetectable ^18^O labels suggested a slow or nonexistent dynamic turnover. At present, we have not fully assigned the identity of the InsP_4_ isomers. They do, however, represent a good example illustrating the potential of rapid ^18^O labeling for dissecting the dynamics of different InsPs pools. Combining the rapid pulse labeling by ^18^O-water with CE-MS allows to generate kinetic data of significantly better time and isomer resolution than the traditional radioactive approaches (Wilson & Saiardi, 2017). Combining this approach with controlled changes in growth conditions or the acute stimulation or inhibition of signaling pathways should allow to obtain novel insights into the regulation and signaling properties of inositol phosphate pools. It will help us to create metabolic models of inositol phosphate-based signaling pathways and to better understand effects exerted by kinase inhibitors on cellular inositol phosphate fluxes, an area of growing interest in e.g. metabolic diseases (Tu-Sekine & Kim, 2022).

In striking contrast to the situation in yeast and mammalian cells we witnessed a sluggish PP-InsP turnover but unexpectedly high turnover of InsP_6_, in the amoeba *Dictyostelium discoideum*. Its lethargic PP-InsPs turnover suggests that the amoeba employs these metabolites differently than yeast or mammalian cells. This correlates with the diversification of the InsP-kinase families that are used in different eukaryotic clades to drive InsPs metabolism and signaling (Laha *et al*, 2022), and with the complexity of the enzymology of PP-InsPs metabolism in amoeba, which is far greater than in yeast or mammalian cells (Desfougères Y *et al*, 2022). The strikingly different behavior of *Dictyostelium* calls for a detailed analysis of InsP fluxes, which may uncover new strategies of InsP signaling. The methods presented here open the door to such investigations.

## Material and methods

### Cell strains and culture media

#### Yeast

The *Saccharomyces cerevisiae* strain BY4741 was used in this study. Yeast cells were shaken (150 rpm, 30°C, unless stated otherwise) in synthetic complete (SC) medium: 6.7 g/l yeast nitrogen base (Formedium, USA) and 2% of glucose.

Medium for ^18^O labeling was prepared as a 2x concentrated SC stock made with normal water in advance, which was then diluted by an equal volume of 99% ^18^O-water (Medical Isotopes Inc) to make SC with 50% ^18^O-water.

#### Mammalian cells

HCT116^UCL^ cells were used in this study. Cells were grown for 6 passages in DMEM-HAM’s F12 (Gibco) supplemented with 5% (v/v) fetal calf serum, 50 U/ml penicillin, 50 mg/ml streptomycin, 5 µg/ml insulin, 5 µg/ml transferrin, and 5 ng/ml selenium. Cells were grown at 37 °C in 5% CO_2_ with 98% humidity.

For ^18^O labeling, a 2x concentrated medium was prepared using the DMEM powder (Sigma) supplemented with 7.4 g/L sodium bicarbonate. This stock was diluted by an equal volume of ^18^O-labeled water to yield the medium with 50% ^18^O-water that was used for the experiments.

#### Dictyostelium

The *Dictyostelium discoideum* strain AX2 obtained from dictyBase (http://dictybase.org) was used in this study. Amoeba were grown at 22°C with gentle shaking (100 rpm) in synthetic SIH medium (Formedium, #SIH0101). Medium for ^18^O labeling was prepared as a 2x concentrated SIH stock made with ddH_2_O, which was then diluted by an equal volume of 99% ^18^O-water (Medical Isotopes Inc) to make SIH with 50% ^18^O-water.

### Extraction of InsPs and ATP

#### Yeast

Yeast cells were grown at 20°C overnight in 50 ml of SC medium to reach logarithmic phase (4.3 x 10^7^ cells/ml) in the morning. 4 ml samples were harvested by centrifugation (3,200 g, 3 min, 20°C) and resuspended in the same volume of SC prepared with 50% of ^18^O-labeled water. Cells were further incubated under the same conditions.

3 ml of yeast culture (4.3 x 10^7^ cells/ml) were mixed with 300 µl of 11 M perchloric acid to a final concentration of 1 M. Samples were snap-frozen in liquid nitrogen and then centrifuged at 20,000x g for 3 min at 4 °C. The soluble supernatant was transferred to a new tube and each sample was supplemented with 6 mg of titanium dioxide (TiO_2_) beads (GL Sciences, Japan), which had been pre-rinsed through two rounds of washing with 1ml of H_2_O and 1 M perchloric acid, respectively. The mixture was gently rotated for 15 min at 4 °C and centrifuged at 20,000x g for 3 min at 4 °C. The TiO_2_ beads were washed twice using 500 µl of 1 M perchloric acid. InsPs and ATP were eluted by incubating the beads with 300 µl of 3% (v/v) NH_4_OH for 5 min at room temperature under gentle shaking. After centrifugation at 20,000x g for 3 min, the eluents were transferred to a new tube. Any remaining TiO_2_ beads were removed by centrifugation at 20,000 g for 3 min. The resulting supernatant was completely dried in a SpeedVac (Labogene, Denmark) at 42 °C. Samples were kept at -20 °C until analysis.

#### Human cells

HCT116^UCL^ cells were seeded in 10 cm^2^ petri dishes and grown to 80% confluence (6.5 x 10^6^ cells) at 37 °C as described above. Cells were detached from the petri dish by adding 5ml of 0.25% Trypsin (Thermo Fisher Scientific, USA) and harvested by centrifugation at 3,200 g for 5 min. The cell pellet was resuspended in 1 ml of DMEM medium (Thermo Fisher Scientific, USA) prepared with 50% of ^18^O-labeled water and further incubated at 37 °C.

At different time points, 1 ml samples were mixed with 100 µl 11 M perchloric acid. After snap-freezing in liquid nitrogen, samples were centrifuged at 16,000 x g in a tabletop centrifuge. The soluble supernatant was transferred into a new tube and mixed with pre-washed TiO_2_ beads (5 mg beads per sample). The extraction was performed in the same way for yeast.

#### Dictyostelium discoideum

*Dictyostelium discoideum* cells were seeded at 2-5 x10^5^ cells/ml and grown for 24-48 hours in 50 ml of SIH medium to reach a cell density of 1-3×10^6^. Amoeba (8×10^6^ per experimental point) were transferred into a 15 ml falcon tube, harvested by centrifugation (800 g, 5 min, 22°C), and resuspended in 2 ml SIH medium prepared with 50% of ^18^O-water. Cells were further incubated under the same conditions. At specific time points, the falcon tube was spun (1000 g, 2 min, 4°C) and the cell pellet snap-frozen in liquid nitrogen. At the end of the experiment, each cell pellet was resuspended in 1 M perchloric acid with 5 mM EDTA and InsPs subjected to TiO_2_ purification as previously described (Wilson *et al*, 2015; Wilson & Saiardi, 2018).

### CE-ESI-MS analysis of InsPs, PP-InsPs and ATP

#### CE-qTOF

An Agilent 7100 capillary electrophoresis system coupled to a qTOF (Agilent 6520) equipped with a commercial CE-MS adapter and sprayer kit (from Agilent) was used. The sheath liquid (water: isopropanol=1:1, v/v) spiked with mass references (TFA anion, [M-H]^-^, 112.9855; HP-0921, [M-H+CH_3_COOH]^-^, 980.0163) was introduced with a constant flow of 6 µl/min. A bare fused silica capillary with a length of 100 cm (50 μm internal diameter and 365 μm outer diameter) was used for CE separation. The background electrolyte (35 mM ammonium acetate titrated with ammonium hydroxide to pH 9.75) was employed for all the experiments. Before the run, the capillary was flushed with background electrolytes for 400 s. 30 or 40 nl of sample was injected by applying pressure (100 mbar for 15 s or 20s). The qTOF was conducted in the negative ionization mode. MS source and scan parameters shown in Supplementary Table S1. Biological samples were analyzed by Acquisition mode MS1. For the ESI-MS fragmentations of ATP and 5-InsP_7_, Acquisition mode Auto MS2 was employed.

#### CE-QQQ

We used an Agilent 7100 capillary electrophoresis system coupled to Agilent 6495C Triple Quadrupole, adopting an Agilent CE-MS interface. The sheath liquid (water: isopropanol=1:1, v/v) was introduced with a constant flow of 10 µl/min. A bare fused silica capillary with a length of 100 cm (50 μm internal diameter and 365 μm outer diameter) was used for CE separation. The background electrolytes (35 mM ammonium acetate titrated with ammonium hydroxide to pH 9.75) was employed for all the experiments. Before running the samples, the capillary was flushed with background electrolytes for 400 s. 30 nl of sample was injected by applying pressure (100 mbar for 15 s). Negative ionization mode was employed. MS source parameters is shown in Supplementary Table S2. For yeast and HCT116 cell samples, MRM transitions settings are shown in Supplementary Table S3, S4, and S5. For amoeba samples, MRM transitions settings are shown in Supplementary Table S3, S6, S7, and S8. All the samples are measured by CE-QQQ unless specifically stated otherwise.

## Conflicts of interest

The authors declare no conflicts of interest.

## Acknowledgments

This study was supported by the Deutsche Forschungsgemeinschaft (DFG) under Germany’s excellence strategy (CIBSS, EXC-2189, Project ID 390939984, to H. J. J., R. G., W. D.). This project has received funding from the European Research Council (ERC) under the European Union’s Horizon 2020 research and innovation program (grant agreement no. 864246, to H. J. J.). H.J. acknowledges funding from the Volkswagen Foundation (VW Momentum Grant 98604). This work was supported by grants from the SNSF (320030-228119, 31003A_179306 and 310030_204713) and ERC (788442) to A. M., and by the HFSP (LT000588/2019) to G. K.

## Notes

### Competing Interest Statement

The authors have declared no competing interest.

## References

Albert C, Safrany TS, Bembenek EM, Reddy KM, Reddy KK, Falck JR, Bröcker M, Shears BS & Mayr WG (1997) Biological variability in the structures of diphosphoinositol polyphosphates in Dictyostelium discoideum and mammalian cells. Biochem J 327: 553– 560

Auesukaree C, Homma T, Tochio H, Shirakawa M, Kaneko Y & Harashima S (2004) Intracellular phosphate serves as a signal for the regulation of the PHO pathway in Saccharomyces cerevisiae. J Biol Chem 279: 17289–17294

Austin S & Mayer A (2020) Phosphate Homeostasis - A Vital Metabolic Equilibrium Maintained Through the INPHORS Signaling Pathway. Front Microbiol 11: 1367

Barker CJ, Wright J, Hughes PJ, Kirk CJ & Michell RH (2004) Complex changes in cellular inositol phosphate complement accompany transit through the cell cycle. Biochem J 380: 465–473

Barneda D, Janardan V, Niewczas I, Collins DM, Cosulich S, Clark J, Stephens LR & Hawkins PT (2022) Acyl chain selection couples the consumption and synthesis of phosphoinositides. Embo J 41: e110038

Berridge MJ & Irvine RF (1984) Inositol trisphosphate, a novel second messenger in cellular signal transduction. Nature 312: 315–321

Blind RD (2020) Structural analyses of inositol phosphate second messengers bound to signaling effector proteins. Adv Biol Regul 75: 100667

Chabert V, Kim G-D, Qiu D, Liu G, Mayer LM, Jamsheer KM, Jessen HJ & Mayer A (2023) Inositol Pyrophosphate Dynamics Reveals Control of the Yeast Phosphate Starvation Program Through 1,5-IP8 and the SPX Domain of Pho81. eLife

Dawis SM, Walseth TF, Deeg MA, Heyman RA, Graeff RM & Goldberg ND (1989) Adenosine triphosphate utilization rates and metabolic pool sizes in intact cells measured by transfer of 18O from water. Biophys J 55: 79–99

Desfougères Y, Portela-Torres P, Qiu D, Livermore TM, Harmel RK, Borghi F, Jessen HJ, Fiedler D & Saiardi A (2022) The inositol pyrophosphate metabolism of Dictyostelium discoideum does not regulate inorganic polyphosphate (polyP) synthesis. Adv Biol Regul 83: 100835

Desfougères Y, Wilson MSC, Laha D, Miller GJ & Saiardi A (2019) ITPK1 mediates the lipid-independent synthesis of inositol phosphates controlled by metabolism. Proc Natl Acad Sci USA 93: 201911431

Dong J, Ma G, Sui L, Wei M, Satheesh V, Zhang R, Ge S, Li J, Zhang T-E, Wittwer C, et al (2019) Inositol Pyrophosphate InsP8 Acts as an Intracellular Phosphate Signal in Arabidopsis. Mol Plant 12: 1463–1473

Dyson JM, Fedele CG, Davies EM, Becanovic J & Mitchell CA (2012) Phosphoinositides I: Enzymes of Synthesis and Degradation. Subcell Biochem 58: 215–279

Eisenbeis VB, Qiu D, Gorka O, Strotmann L, Liu G, Prucker I, Su XB, Wilson MSC, Ritter K, Loenarz C, et al (2023) β-lapachone regulates mammalian inositol pyrophosphate levels in an NQO1- and oxygen-dependent manner. Proc Natl Acad Sci 120: e2306868120

Falkenburger BH, Jensen JB, Dickson EJ, Suh B & Hille B (2010) SYMPOSIUM REVIEW: Phosphoinositides: lipid regulators of membrane proteins. J Physiology 588: 3179–3185

Goldbeter A & Koshland DE (1981) An amplified sensitivity arising from covalent modification in biological systems. Proc Natl Acad Sci 78: 6840–6844

Goldbeter A & Koshland DE (1987) Energy expenditure in the control of biochemical systems by covalent modification. J Biol Chem 262: 4460–71

Gu C, Liu J, Liu X, Zhang H, Luo J, Wang H, Locasale JW & Shears SB (2021) Metabolic supervision by PPIP5K, an inositol pyrophosphate kinase/phosphatase, controls proliferation of the HCT116 tumor cell line. Proc Natl Acad Sci USA 118

Haas TM, Mundinger S, Qiu D, Jork N, Ritter K, Dürr-Mayer T, Ripp A, Saiardi A, Schaaf G & Jessen HJ (2021) Stable isotope phosphate labelling of diverse metabolites is enabled by a family of 18O-phosphoramidites. Angewandte Chemie Int Ed

Hackney DD (1980) Theoretical analysis of distribution of [18O]Pi species during exchange with water. Application to exchanges catalyzed by yeast inorganic pyrophosphatase. J Biological Chem 255: 5320–8

Harmel RK, Puschmann R, Trung MN, Saiardi A, Schmieder P & Fiedler D (2019) Harnessing 13C-labeled myo-inositol to interrogate inositol phosphate messengers by NMR. Chem Sci 10: 5267–5274

Heijnen JJ (2010) Biosystems Engineering II, Linking Cellular Networks and Bioprocesses. Adv Biochem Eng Biotechnology 121: 139–162

Hofer A, Cremosnik GS, Müller AC, Giambruno R, Trefzer C, Superti-Furga G, Bennett KL & Jessen HJ (2015) A Modular Synthesis of Modified Phosphoanhydrides. Chem European J 21: 10116–10122

Juranić N, Nemutlu E, Zhang S, Dzeja P, Terzic A & Macura S (2011) 31P NMR correlation maps of 18O/16O chemical shift isotopic effects for phosphometabolite labeling studies. J Biomol Nmr 50: 237–245

Kenyon CP, Roth RL, Westhuyzen CW van der & Parkinson CJ (2012) Conserved phosphoryl transfer mechanisms within kinase families and the role of the C8 proton of ATP in the activation of phosphoryl transfer. BMC Res Notes 5: 131

Kochanowski K, Volkmer B, Gerosa L, Rijsewijk BRH van, Schmidt A & Heinemann M (2013) Functioning of a metabolic flux sensor in Escherichia coli. Proc Natl Acad Sci 110: 1130–1135

Kotte O, Zaugg JB & Heinemann M (2010) Bacterial adaptation through distributed sensing of metabolic fluxes. Mol Syst Biol 6: 355–355

Lassila JK, Zalatan JG & Herschlag D (2011) Biological Phosphoryl-Transfer Reactions: Understanding Mechanism and Catalysis. Annu Rev Biochem 80: 669–702

Lee W-K, Son SH, Jin B-S, Na J-H, Kim S-Y, Kim K-H, Kim EE, Yu YG & Lee HH (2013) Structural and functional insights into the regulation mechanism of CK2 by IP6 and the intrinsically disordered protein Nopp140. Proc Natl Acad Sci USA 110: 19360–19365

Li X, Gu C, Hostachy S, Sahu S, Wittwer C, Jessen HJ, Fiedler D, Wang H & Shears SB (2020) Control of XPR1-dependent cellular phosphate efflux by InsP8 is an exemplar for functionally-exclusive inositol pyrophosphate signaling. Proc Natl Acad Sci USA 14: 201908830

Li Y, Cross FR & Chait BT (2014) Method for identifying phosphorylated substrates of specific cyclin/cyclin-dependent kinase complexes. Proc National Acad Sci 111: 11323– 11328

Liu G, Riemer E, Schneider R, Cabuzu D, Bonny O, Wagner CA, Qiu D, Saiardi A, Strauss A, Lahaye T, et al (2023) The phytase RipBL1 enables the assignment of a specific inositol phosphate isomer as a structural component of human kidney stones. RSC Chem Biol 4: 300–309

Macbeth MR, Schubert HL, VanDemark AP, Lingam AT, Hill CP & Bass BL (2005) Inositol Hexakisphosphate Is Bound in the ADAR2 Core and Required for RNA Editing. Science 309: 1534–1539

Maffucci T & Falasca M (2020) Signalling Properties of Inositol Polyphosphates. Molecules 25: 5281

Menniti FS, Miller RN, Putney JW & Shears SB (1993) Turnover of inositol polyphosphate pyrophosphates in pancreatoma cells. J Biol Chem 268: 3850–3856

Miller CA & Beard DA (2008) The Effects of Reversibility and Noise on Stochastic Phosphorylation Cycles and Cascades. Biophys J 95: 2183–2192

Mitchell RA, Lamos CM & Russo JA (1980) Mitochondrial-catalyzed ATP hydrolysis in highly enriched [18O]H2O. Frequency distributions of 18O-labelled Pi species. Biochim Biophys Acta (BBA) - Bioenerg 592: 406–414

Monge ME, Dodds JN, Baker ES, Edison AS & Fernández FM (2019) Challenges in Identifying the Dark Molecules of Life. Annu Rev Anal Chem 12: 1–23

Monserrate JP & York JD (2010) Inositol phosphate synthesis and the nuclear processes they affect. Curr Opin Cell Biol 22: 365–373

Moritoh Y, Abe S-I, Akiyama H, Kobayashi A, Koyama R, Hara R, Kasai S & Watanabe M (2021) The enzymatic activity of inositol hexakisphosphate kinase controls circulating phosphate in mammals. Nature communications 12: 4847–15

Müller AC, Giambruno R, Weißer J, Májek P, Hofer A, Bigenzahn JW, Superti-Furga G, Jessen HJ & Bennett KL (2016) Identifying Kinase Substrates via a Heavy ATP Kinase Assay and Quantitative Mass Spectrometry. Sci Rep-uk 6: 28107

Nemutlu E, Gupta A, Zhang S, Viqar M, Holmuhamedov E, Terzic A, Jahangir A & Dzeja P (2015) Decline of Phosphotransfer and Substrate Supply Metabolic Circuits Hinders ATP Cycling in Aging Myocardium. PLoS ONE 10: e0136556

Nemutlu E, Juranic N, Zhang S, Ward LE, Dutta T, Nair KS, Terzic A, Macura S & Dzeja PP (2012) Electron spray ionization mass spectrometry and 2D 31P NMR for monitoring 18O/16O isotope exchange and turnover rates of metabolic oligophosphates. Anal Bioanal Chem 403: 697–706

Obr M, Schur FKM & Dick RA (2021) A Structural Perspective of the Role of IP6 in Immature and Mature Retroviral Assembly. Viruses 13: 1853

Pisani F, Livermore T, Rose G, Chubb JR, Gaspari M & Saiardi A (2014) Analysis of Dictyostelium discoideum Inositol Pyrophosphate Metabolism by Gel Electrophoresis. PLoS ONE 9: e85533

Posor Y, Jang W & Haucke V (2022) Phosphoinositides as membrane organizers. Nat Rev Mol Cell Bio 23: 797–816

Qiu D, Gu C, Liu G, Ritter K, Eisenbeis VB, Bittner T, Gruzdev A, Seidel L, Bengsch B, Shears SB, et al (2022) Capillary electrophoresis mass spectrometry identifies new isomers of inositol pyrophosphates in mammalian tissues. Chem Sci 14: 658–667

Qiu D, Wilson MS, Eisenbeis VB, Harmel RK, Riemer E, Haas TM, Wittwer C, Jork N, Gu C, Shears SB, et al (2020) Analysis of inositol phosphate metabolism by capillary electrophoresis electrospray ionization mass spectrometry. Nature communications 11: 6035–12

Raser JM & O’Shea EK (2004) Control of stochasticity in eukaryotic gene expression. Science 304: 1811–1814

Renner N, Kleinpeter A, Mallery DL, Albecka A, Faysal KMR, Böcking T, Saiardi A, Freed EO & James LC (2023) HIV-1 is dependent on its immature lattice to recruit IP6 for mature capsid assembly. Nat Struct Mol Biol 30: 370–382

Riemer E, Qiu D, Laha D, Harmel RK, Gaugler P, Gaugler V, Frei M, Hajirezaei M-R, Laha NP, Krusenbaum L, et al (2021) ITPK1 is an InsP6/ADP phosphotransferase that controls phosphate signaling in Arabidopsis. Mol Plant

Sahu S, Gordon J, Gu C, Sobhany M, Fiedler D, Stanley RE & Shears SB (2023) Nucleolar Architecture Is Modulated by a Small Molecule, the Inositol Pyrophosphate 5-InsP7. Biomolecules 13: 153

Samoilov M, Plyasunov S & Arkin AP (2005) Stochastic amplification and signaling in enzymatic futile cycles through noise-induced bistability with oscillations. Proc National Acad Sci 102: 2310–2315

Shears SB, Ganapathi SB, Gokhale NA, Schenk TMH, Wang H, Weaver JD, Zaremba A & Zhou Y (2012) Phosphoinositides II: The Diverse Biological Functions. Subcell Biochem 59: 389–412

Szijgyarto Z, Garedew A, Azevedo C & Saiardi A (2011) Influence of inositol pyrophosphates on cellular energy dynamics. Science 334: 802–805

Trung MN, Kieninger S, Fandi Z, Qiu D, Liu G, Mehendale NK, Saiardi A, Jessen H, Keller B & Fiedler D (2022) Stable Isotopomers of myo-Inositol Uncover a Complex MINPP1-Dependent Inositol Phosphate Network. Acs Central Sci 8: 1683–1694

Tsui MM & York JD (2010) Roles of inositol phosphates and inositol pyrophosphates in development, cell signaling and nuclear processes. Adv Enzym Regul 50: 324–337

Tu-Sekine B & Kim SF (2022) The Inositol Phosphate System—A Coordinator of Metabolic Adaptability. Int J Mol Sci 23: 6747

Vardi N, Levy S, Assaf M, Carmi M & Barkai N (2013) Budding Yeast Escape Commitment to the Phosphate Starvation Program Using Gene Expression Noise. Curr Biol

Vardi N, Levy S, Gurvich Y, Polacheck T, Carmi M, Jaitin D, Amit I & Barkai N (2014) Sequential feedback induction stabilizes the phosphate starvation response in budding yeast. Cell Rep 9: 1122–1134

Versaw WK & Metzenberg RL (1996) Intracellular phosphate--water oxygen exchange measured by mass spectrometry. Anal Biochem 241: 14–17

Walker JE (1998) ATP Synthesis by Rotary Catalysis (Nobel lecture). Angew Chem Int Ed 37: 2308–2319

Wang L, Amelung W & Willbold S (2021) 18O Isotope Labeling Combined with 31P Nuclear Magnetic Resonance Spectroscopy for Accurate Quantification of Hydrolyzable Phosphorus Species in Environmental Samples. Anal Chem 93: 2018–2025

Wild R, Gerasimaite R, Jung J-Y, Truffault V, Pavlovic I, Schmidt A, Saiardi A, Jessen HJ, Poirier Y, Hothorn M, et al (2016) Control of eukaryotic phosphate homeostasis by inositol polyphosphate sensor domains. Science 352: 986–990

Wilson M & Saiardi A (2018) Inositol Phosphates Purification Using Titanium Dioxide Beads. BIO-Protoc 8

Wilson MSC, Bulley SJ, Pisani F, Irvine RF & Saiardi A (2015) A novel method for the purification of inositol phosphates from biological samples reveals that no phytate is present in human plasma or urine. Open Biol 5: 150014

Wilson MSC & Saiardi A (2017) Importance of Radioactive Labelling to Elucidate Inositol Polyphosphate Signalling. Top Curr Chem (Cham) 375: 14

Wykoff DD, Rizvi AH, Raser JM, Margolin B & O’Shea EK (2007) Positive feedback regulates switching of phosphate transporters in S. cerevisiae. Mol Cell 27: 1005–1013

