## supplement for "Pools of independently cycling inositol phosphates revealed by pulse labeling with ^18^O-water"

### Supplementary Information

Supplementary Table S1. Instrument and scan source parameters of qTOF

| **Instrument and scan source parameters** | |
|---|---|
| Gas Temperature | 250 °C |
| Gas Flow | 3 L/min |
| Nebulizer | 10 psi |
| Vcap | 3500 V |
| Fragmentor | 100 V |
| Skimmer | 65 V |
| OctopoleRFPeak | 750 V |
| **Acquisition mode MS1** | |
| Scan *m/z* Range | 80-1500 |
| Scan Rate | 1 spectra/sec |
| **Acquisition mode AutoMS2** | |
| Scan *m/z* Range | 100-1000 |
| Scan Rate | 1 spectra/sec |
| Isolation Width MS/MS | Medium (˜4 amu) |
| Using Fixed Collision Energies | 5 V, 10 V, 20 V, 30 V, 40V |
| Max Precursors per Cycle | 2 |
| Threshold | 200 Abs |
| Threshold (Rel) | 0.01 |
| Sort Precursors | By abundance only |
| Charge State Preference | 1, 2 |

Supplementary Table S2. Instrument and scan source parameters of QQQ

| **Source Parameters** | |
|---|---|
| Gas Temperature | 150 °C |
| Gas Flow | 11 L/min |
| Nebulizer | 8 psi |
| Sheath Gas Temperature | 175 °C |
| Sheath Gas Flow | 8 L/min |
| Capillary Voltage | -2000 V |
| Nozzle Voltage | 2000 V |
| High Pressure RF (Ion Funnel Parameters) | 70 V |
| Low Pressure RF (Ion Funnel Parameters) | 40 V |

Supplementary Table S3. MRM transitions setting of ATP for the measurement of $^{18}O$ labeled samples

| Molecular name | Precursor Ion | Product Ion | Type of transition | Collision Energy (V) | Cell Accelerator Voltage |
|---|---|---|---|---|---|
| [$^{18}O_7$] ATP | 520 | 420 | $[M\text{-}H]^- \rightarrow [M\text{-}H\text{-}H_3PO_3{}^{18}O]^-$ | 25 | 1 |
| [$^{18}O_7$] ATP | 520 | 418 | $[M\text{-}H]^- \rightarrow [M\text{-}H\text{-}H_3PO_2{}^{18}O_2]^-$ | 25 | 1 |
| [$^{18}O_7$] ATP | 520 | 416 | $[M\text{-}H]^- \rightarrow [M\text{-}H\text{-}H_3PO{}^{18}O_3]^-$ | 25 | 1 |
| [$^{18}O_7$] ATP | 520 | 414 | $[M\text{-}H]^- \rightarrow [M\text{-}H\text{-}H_3P{}^{18}O_4]^-$ | 25 | 1 |
| [$^{18}O_6$] ATP | 518 | 420 | $[M\text{-}H]^- \rightarrow [M\text{-}H\text{-}H_3PO_4]^-$ | 25 | 1 |
| [$^{18}O_6$] ATP | 518 | 418 | $[M\text{-}H]^- \rightarrow [M\text{-}H\text{-}H_3PO_3{}^{18}O]^-$ | 25 | 1 |
| [$^{18}O_6$] ATP | 518 | 416 | $[M\text{-}H]^- \rightarrow [M\text{-}H\text{-}H_3PO_2{}^{18}O_2]^-$ | 25 | 1 |
| [$^{18}O_6$] ATP | 518 | 414 | $[M\text{-}H]^- \rightarrow [M\text{-}H\text{-}H_3PO{}^{18}O_3]^-$ | 25 | 1 |
| [$^{18}O_6$] ATP | 518 | 412 | $[M\text{-}H]^- \rightarrow [M\text{-}H\text{-}H_3P{}^{18}O_4]^-$ | 25 | 1 |
| [$^{18}O_5$] ATP | 516 | 416 | $[M\text{-}H]^- \rightarrow [M\text{-}H\text{-}H_3PO_3{}^{18}O]^-$ | 25 | 1 |
| [$^{18}O_5$] ATP | 516 | 414 | $[M\text{-}H]^- \rightarrow [M\text{-}H\text{-}H_3PO_2{}^{18}O_2]^-$ | 25 | 1 |
| [$^{18}O_5$] ATP | 516 | 412 | $[M\text{-}H]^- \rightarrow [M\text{-}H\text{-}H_3PO{}^{18}O_3]^-$ | 25 | 1 |
| [$^{18}O_5$] ATP | 516 | 410 | $[M\text{-}H]^- \rightarrow [M\text{-}H\text{-}H_3P{}^{18}O_4]^-$ | 25 | 1 |
| [$^{18}O_4$] ATP | 514 | 416 | $[M\text{-}H]^- \rightarrow [M\text{-}H\text{-}H_3PO_4]^-$ | 25 | 1 |
| [$^{18}O_4$] ATP | 514 | 414 | $[M\text{-}H]^- \rightarrow [M\text{-}H\text{-}H_3PO_3{}^{18}O]^-$ | 25 | 1 |
| [$^{18}O_4$] ATP | 514 | 412 | $[M\text{-}H]^- \rightarrow [M\text{-}H\text{-}H_3PO_2{}^{18}O_2]^-$ | 25 | 1 |
| [$^{18}O_4$] ATP | 514 | 410 | $[M\text{-}H]^- \rightarrow [M\text{-}H\text{-}H_3PO{}^{18}O_3]^-$ | 25 | 1 |
| [$^{18}O_4$] ATP | 514 | 408 | $[M\text{-}H]^- \rightarrow [M\text{-}H\text{-}H_3P{}^{18}O_4]^-$ | 25 | 1 |
| [$^{18}O_3$] ATP | 512 | 414 | $[M\text{-}H]^- \rightarrow [M\text{-}H\text{-}H_3PO_4]^-$ | 25 | 1 |
| [$^{18}O_3$] ATP | 512 | 412 | $[M\text{-}H]^- \rightarrow [M\text{-}H\text{-}H_3PO_3{}^{18}O]^-$ | 25 | 1 |
| [$^{18}O_3$] ATP | 512 | 410 | $[M\text{-}H]^- \rightarrow [M\text{-}H\text{-}H_3PO_2{}^{18}O_2]^-$ | 25 | 1 |
| [$^{18}O_3$] ATP | 512 | 408 | $[M\text{-}H]^- \rightarrow [M\text{-}H\text{-}H_3PO{}^{18}O_3]^-$ | 25 | 1 |
| [$^{18}O_2$] ATP | 510 | 412 | $[M\text{-}H]^- \rightarrow [M\text{-}H\text{-}H_3PO_4]^-$ | 25 | 1 |
| [$^{18}O_2$] ATP | 510 | 410 | $[M\text{-}H]^- \rightarrow [M\text{-}H\text{-}H_3PO_3{}^{18}O]^-$ | 25 | 1 |
| [$^{18}O_2$] ATP | 510 | 408 | $[M\text{-}H]^- \rightarrow [M\text{-}H\text{-}H_3PO_2{}^{18}O_2]^-$ | 25 | 1 |
| [$^{18}O$] ATP | 508 | 410 | $[M\text{-}H]^- \rightarrow [M\text{-}H\text{-}H_3PO_3{}^{18}O]^-$ | 25 | 1 |
| [$^{18}O$] ATP | 508 | 408 | $[M\text{-}H]^- \rightarrow [M\text{-}H\text{-}H_3PO_2{}^{18}O_2]^-$ | 25 | 1 |
| ATP | 506 | 408 | $[M\text{-}H]^- \rightarrow [M\text{-}H\text{-}H_3PO_4]^-$ | 25 | 1 |
| [$^{13}C_{10}$] ATP | 516 | 418 | $[M\text{-}H]^- \rightarrow [M\text{-}H\text{-}H_3PO_4]^-$ | 25 | 1 |

Notes: Negative polarity; fragmentor (V) is 166; dwell is 40.

Supplementary Table S4. MRM transitions setting of $InsP_{6-8}$ for the measurement of yeast and human cells samples

| Molecular name | Precursor Ion | Product Ion | Type of transition | Collision Energy (V) | Cell Accelerator Voltage |
|---|---|---|---|---|---|
| $[^{13}C_6]$ $InsP_8$ | 411.9 | 362.9 | $[M\text{-}2H]^{2-} \rightarrow [M\text{-}2H\text{-}H_3PO_4]^{2-}$ | 10 | 1 |
| $InsP_8$ | 408.9 | 359.9 | $[M\text{-}2H]^{2-} \rightarrow [M\text{-}2H\text{-}H_3PO_4]^{2-}$ | 10 | 1 |
| $[^{18}O]$ $InsP_8$ | 409.9 | 360.9 | $[M\text{-}2H]^{2-} \rightarrow [M\text{-}2H\text{-}H_3PO_4]^{2-}$ | 10 | 1 |
| $[^{18}O]$ $InsP_8$ | 409.9 | 359.9 | $[M\text{-}2H]^{2-} \rightarrow [M\text{-}2H\text{-}H_3PO_3{}^{18}O]^{2-}$ | 10 | 1 |
| $[^{18}O_2]$ $InsP_8$ | 410.9 | 361.9 | $[M\text{-}2H]^{2-} \rightarrow [M\text{-}2H\text{-}H_3PO_4]^{2-}$ | 10 | 1 |
| $[^{18}O_2]$ $InsP_8$ | 410.9 | 360.9 | $[M\text{-}2H]^{2-} \rightarrow [M\text{-}2H\text{-}H_3PO_3{}^{18}O]^{2-}$ | 10 | 1 |
| $[^{18}O_2]$ $InsP_8$ | 410.9 | 359.9 | $[M\text{-}2H]^{2-} \rightarrow [M\text{-}2H\text{-}H_3PO_2{}^{18}O_2]^{2-}$ | 10 | 1 |
| $[^{13}C_6]$ $InsP_7$ | 371.9 | 322.9 | $[M\text{-}2H]^{2-} \rightarrow [M\text{-}2H\text{-}H_3PO_4]^{2-}$ | 10 | 3 |
| $InsP_7$ | 368.9 | 319.9 | $[M\text{-}2H]^{2-} \rightarrow [M\text{-}2H\text{-}H_3PO_4]^{2-}$ | 10 | 3 |
| $[^{18}O]$ $InsP_7$ | 369.9 | 320.9 | $[M\text{-}2H]^{2-} \rightarrow [M\text{-}2H\text{-}H_3PO_4]^{2-}$ | 10 | 3 |
| $[^{18}O]$ $InsP_7$ | 369.9 | 319.9 | $[M\text{-}2H]^{2-} \rightarrow [M\text{-}2H\text{-}H_3PO_3{}^{18}O]^{2-}$ | 10 | 3 |
| $[^{18}O_2]$ $InsP_7$ | 370.9 | 321.9 | $[M\text{-}2H]^{2-} \rightarrow [M\text{-}2H\text{-}H_3PO_4]^{2-}$ | 10 | 3 |
| $[^{18}O_2]$ $InsP_7$ | 370.9 | 320.9 | $[M\text{-}2H]^{2-} \rightarrow [M\text{-}2H\text{-}H_3PO_3{}^{18}O]^{2-}$ | 10 | 3 |
| $[^{18}O_2]$ $InsP_7$ | 370.9 | 319.9 | $[M\text{-}2H]^{2-} \rightarrow [M\text{-}2H\text{-}H_3PO_2{}^{18}O_2]^{2-}$ | 10 | 3 |
| $[^{13}C_6]$ $InsP_6$ | 331.9 | 486.9 | $[M\text{-}2H]^{2-} \rightarrow [M\text{-}H\text{-}HPO_3\text{-}H_3PO_4]^{-}$ | 13 | 4 |
| $InsP_6$ | 328.9 | 480.9 | $[M\text{-}2H]^{2-} \rightarrow [M\text{-}H\text{-}HPO_3\text{-}H_3PO_4]^{-}$ | 13 | 4 |
| $[^{18}O]$ $InsP_6$ | 329.9 | 482.9 | $[M\text{-}2H]^{2-} \rightarrow [M\text{-}H\text{-}HPO_3\text{-}H_3PO_4]^{-}$ | 13 | 4 |
| $[^{18}O]$ $InsP_6$ | 329.9 | 480.9 | $[M\text{-}2H]^{2-} \rightarrow [M\text{-}H\text{-}HPO_3\text{-} H_3PO_3{}^{18}O]^{-}$ | 13 | 4 |
| $[^{18}O_2]$ $InsP_6$ | 330.9 | 484.9 | $[M\text{-}2H]^{2-} \rightarrow [M\text{-}H\text{-}HPO_3\text{-}H_3PO_4]^{-}$ | 13 | 4 |
| $[^{18}O_2]$ $InsP_6$ | 330.9 | 482.9 | $[M\text{-}2H]^{2-} \rightarrow [M\text{-}H\text{-}HPO_3\text{-}H_3PO_3{}^{18}O]^{-}$ | 13 | 4 |
| $[^{18}O_2]$ $InsP_6$ | 330.9 | 480.9 | $[M\text{-}2H]^{2-} \rightarrow [M\text{-}H\text{-}HPO_3\text{-}H_3PO_2{}^{18}O_2]^{-}$ | 13 | 4 |

Notes: Negative polarity; fragmentor (V) is 166; dwell is 80.

Supplementary Table S5. MRM transitions setting of $InsP_{3-5}$ for the measurement of yeast and human cells samples

| Molecular name | Precursor Ion | Product Ion | Type of transition | Collision Energy (V) | Cell Accelerator Voltage |
|---|---|---|---|---|---|
| [$^{13}C_6$] $InsP_5$ | 292 | 504.9 | $[M-2H]^{2-} \rightarrow [M-H-HPO_3]^-$ | 9 | 3 |
| $InsP_5$ | 289 | 498.9 | $[M-2H]^{2-} \rightarrow [M-H-HPO_3]^-$ | 9 | 3 |
| [$^{18}O$] $InsP_5$ | 290 | 500.9 | $[M-2H]^{2-} \rightarrow [M-H-HPO_3]^-$ | 9 | 3 |
| [$^{18}O$] $InsP_5$ | 290 | 498.9 | $[M-2H]^{2-} \rightarrow [M-H-HPO_2{}^{18}O]^-$ | 9 | 3 |
| [$^{18}O_2$] $InsP_5$ | 291 | 502.9 | $[M-2H]^{2-} \rightarrow [M-H-HPO_3]^-$ | 9 | 3 |
| [$^{18}O_2$] $InsP_5$ | 291 | 500.9 | $[M-2H]^{2-} \rightarrow [M-H-HPO_2{}^{18}O]^-$ | 9 | 3 |
| [$^{18}O_2$] $InsP_5$ | 291 | 498.9 | $[M-2H]^{2-} \rightarrow [M-H-HPO{}^{18}O_2]^-$ | 9 | 3 |
| $InsP_4$ | 249 | 418.9 | $[M-2H]^{2-} \rightarrow [M-H-HPO_3]^-$ | 5 | 1 |
| [$^{18}O$] $InsP_4$ | 250 | 420.9 | $[M-2H]^{2-} \rightarrow [M-H-HPO_3]^-$ | 10 | 3 |
| [$^{18}O$] $InsP_4$ | 250 | 418.9 | $[M-2H]^{2-} \rightarrow [M-H-HPO_2{}^{18}O]^-$ | 10 | 3 |
| [$^{18}O_2$] $InsP_4$ | 251 | 422.9 | $[M-2H]^{2-} \rightarrow [M-H-HPO_3]^-$ | 10 | 3 |
| [$^{18}O_2$] $InsP_4$ | 251 | 420.9 | $[M-2H]^{2-} \rightarrow [M-H-HPO_2{}^{18}O]^-$ | 10 | 3 |
| [$^{18}O_2$] $InsP_4$ | 251 | 418.9 | $[M-2H]^{2-} \rightarrow [M-H-HPO{}^{18}O_2]^-$ | 10 | 3 |
| $InsP_3$ | 418.9 | 320.8 | $[M-H]^- \rightarrow [M-H-H_3PO_4]^-$ | 17 | 4 |
| [$^{18}O$] $InsP_3$ | 420.9 | 322.8 | $[M-H]^- \rightarrow [M-H-H_3PO_4]^-$ | 17 | 4 |
| [$^{18}O$] $InsP_3$ | 420.9 | 320.8 | $[M-H]^- \rightarrow [M-H-H_3PO_3{}^{18}O]^-$ | 17 | 4 |
| [$^{18}O_2$] $InsP_3$ | 422.9 | 324.9 | $[M-H]^- \rightarrow [M-H-H_3PO_4]^-$ | 17 | 4 |
| [$^{18}O_2$] $InsP_3$ | 422.9 | 322.9 | $[M-H]^- \rightarrow [M-H-H_3PO_3{}^{18}O]^-$ | 17 | 4 |
| [$^{18}O_2$] $InsP_3$ | 422.9 | 320.9 | $[M-H]^- \rightarrow [M-H-H_3PO_2{}^{18}O_2]^-$ | 17 | 4 |

Notes: Negative polarity; fragmentor (V) is 166; dwell is 60

Supplementary Table S6. MRM transitions setting of $InsP_8$ for amoeba samples

| Molecular name | Precursor Ion | Product Ion | Type of transition | Collision Energy (V) | Cell Accelerator Voltage |
|---|---|---|---|---|---|
| $InsP_8$ | 408.9 | 359.9 | $[M-2H]^{2-} \rightarrow [M-2H-H_3PO_4]^{2-}$ | 10 | 1 |
| $[^{18}O]$ $InsP_8$ | 409.9 | 360.9 | $[M-2H]^{2-} \rightarrow [M-2H-H_3PO_4]^{2-}$ | 10 | 1 |
| $[^{18}O]$ $InsP_8$ | 409.9 | 359.9 | $[M-2H]^{2-} \rightarrow [M-2H-H_3PO_3{}^{18}O]^{2-}$ | 10 | 1 |
| $[^{18}O_2]$ $InsP_8$ | 410.9 | 361.9 | $[M-2H]^{2-} \rightarrow [M-2H-H_3PO_4]^{2-}$ | 10 | 1 |
| $[^{18}O_2]$ $InsP_8$ | 410.9 | 360.9 | $[M-2H]^{2-} \rightarrow [M-2H-H_3PO_3{}^{18}O]^{2-}$ | 10 | 1 |
| $[^{18}O_2]$ $InsP_8$ | 410.9 | 359.9 | $[M-2H]^{2-} \rightarrow [M-2H-H_3PO_2{}^{18}O_2]^{2-}$ | 10 | 1 |
| $[^{18}O_3]$ $InsP_8$ | 411.9 | 362.9 | $[M-2H]^{2-} \rightarrow [M-2H-H_3PO_4]^{2-}$ | 10 | 1 |
| $[^{18}O_3]$ $InsP_8$ | 411.9 | 361.9 | $[M-2H]^{2-} \rightarrow [M-2H-H_3PO_3{}^{18}O]^{2-}$ | 10 | 1 |
| $[^{18}O_3]$ $InsP_8$ | 411.9 | 360.9 | $[M-2H]^{2-} \rightarrow [M-2H-H_3PO_2{}^{18}O_2]^{2-}$ | 10 | 1 |
| $[^{18}O_3]$ $InsP_8$ | 411.9 | 359.9 | $[M-2H]^{2-} \rightarrow [M-2H-H_3PO{}^{18}O_3]^{2-}$ | 10 | 1 |
| $[^{18}O_4]$ $InsP_8$ | 412.9 | 363.9 | $[M-2H]^{2-} \rightarrow [M-2H-H_3PO_4]^{2-}$ | 10 | 1 |
| $[^{18}O_4]$ $InsP_8$ | 412.9 | 362.9 | $[M-2H]^{2-} \rightarrow [M-2H-H_3PO_3{}^{18}O]^{2-}$ | 10 | 1 |
| $[^{18}O_4]$ $InsP_8$ | 412.9 | 361.9 | $[M-2H]^{2-} \rightarrow [M-2H-H_3PO_2{}^{18}O_2]^{2-}$ | 10 | 1 |
| $[^{18}O_4]$ $InsP_8$ | 412.9 | 360.9 | $[M-2H]^{2-} \rightarrow [M-2H-H_3PO{}^{18}O_3]^{2-}$ | 10 | 1 |
| $[^{18}O_4]$ $InsP_8$ | 412.9 | 359.9 | $[M-2H]^{2-} \rightarrow [M-2H-H_3P{}^{18}O_4]^{2-}$ | 10 | 1 |
| $[^{18}O_5]$ $InsP_8$ | 413.9 | 364.9 | $[M-2H]^{2-} \rightarrow [M-2H-H_3PO_4]^{2-}$ | 10 | 1 |
| $[^{18}O_5]$ $InsP_8$ | 413.9 | 363.9 | $[M-2H]^{2-} \rightarrow [M-2H-H_3PO_3{}^{18}O]^{2-}$ | 10 | 1 |
| $[^{18}O_5]$ $InsP_8$ | 413.9 | 362.9 | $[M-2H]^{2-} \rightarrow [M-2H-H_3PO_2{}^{18}O_2]^{2-}$ | 10 | 1 |
| $[^{18}O_5]$ $InsP_8$ | 413.9 | 361.9 | $[M-2H]^{2-} \rightarrow [M-2H-H_3PO{}^{18}O_3]^{2-}$ | 10 | 1 |
| $[^{18}O_5]$ $InsP_8$ | 413.9 | 360.9 | $[M-2H]^{2-} \rightarrow [M-2H-H_3P{}^{18}O_4]^{2-}$ | 10 | 1 |
| $[^{18}O_6]$ $InsP_8$ | 414.9 | 365.9 | $[M-2H]^{2-} \rightarrow [M-2H-H_3PO_4]^{2-}$ | 10 | 1 |
| $[^{18}O_6]$ $InsP_8$ | 414.9 | 364.9 | $[M-2H]^{2-} \rightarrow [M-2H-H_3PO_3{}^{18}O]^{2-}$ | 10 | 1 |
| $[^{18}O_6]$ $InsP_8$ | 414.9 | 363.9 | $[M-2H]^{2-} \rightarrow [M-2H-H_3PO_2{}^{18}O_2]^{2-}$ | 10 | 1 |
| $[^{18}O_6]$ $InsP_8$ | 414.9 | 362.9 | $[M-2H]^{2-} \rightarrow [M-2H-H_3PO{}^{18}O_3]^{2-}$ | 10 | 1 |
| $[^{18}O_6]$ $InsP_8$ | 414.9 | 361.9 | $[M-2H]^{2-} \rightarrow [M-2H-H_3P{}^{18}O_4]^{2-}$ | 10 | 1 |

Notes: Negative polarity; fragmentor (V) is 166; dwell is 25.

Supplementary Table S7. MRM transitions setting of $InsP_7$ for amoeba samples

| Molecular name | Precursor Ion | Product Ion | Type of transition | Collision Energy (V) | Cell Accelerator Voltage |
|---|---|---|---|---|---|
| $InsP_7$ | 368.9 | 319.9 | $[M-2H]^{2-} \rightarrow [M-2H-H_3PO_4]^{2-}$ | 10 | 3 |
| $[^{18}O]$ $InsP_7$ | 369.9 | 320.9 | $[M-2H]^{2-} \rightarrow [M-2H-H_3PO_4]^{2-}$ | 10 | 3 |
| $[^{18}O]$ $InsP_7$ | 369.9 | 319.9 | $[M-2H]^{2-} \rightarrow [M-2H-H_3PO_3{}^{18}O]^{2-}$ | 10 | 3 |
| $[^{18}O_2]$ $InsP_7$ | 370.9 | 321.9 | $[M-2H]^{2-} \rightarrow [M-2H-H_3PO_4]^{2-}$ | 10 | 3 |
| $[^{18}O_2]$ $InsP_7$ | 370.9 | 320.9 | $[M-2H]^{2-} \rightarrow [M-2H-H_3PO_3{}^{18}O]^{2-}$ | 10 | 3 |
| $[^{18}O_2]$ $InsP_7$ | 370.9 | 319.9 | $[M-2H]^{2-} \rightarrow [M-2H-H_3PO_2{}^{18}O_2]^{2-}$ | 10 | 3 |
| $[^{18}O_3]$ $InsP_7$ | 371.9 | 322.9 | $[M-2H]^{2-} \rightarrow [M-2H-H_3PO_4]^{2-}$ | 10 | 3 |
| $[^{18}O_3]$ $InsP_7$ | 371.9 | 321.9 | $[M-2H]^{2-} \rightarrow [M-2H-H_3PO_3{}^{18}O]^{2-}$ | 10 | 3 |
| $[^{18}O_3]$ $InsP_7$ | 371.9 | 320.9 | $[M-2H]^{2-} \rightarrow [M-2H-H_3PO_2{}^{18}O_2]^{2-}$ | 10 | 3 |
| $[^{18}O_3]$ $InsP_7$ | 371.9 | 319.9 | $[M-2H]^{2-} \rightarrow [M-2H-H_3PO{}^{18}O_3]^{2-}$ | 10 | 3 |
| $[^{18}O_4]$ $InsP_7$ | 372.9 | 323.9 | $[M-2H]^{2-} \rightarrow [M-2H-H_3PO_4]^{2-}$ | 10 | 3 |
| $[^{18}O_4]$ $InsP_7$ | 372.9 | 322.9 | $[M-2H]^{2-} \rightarrow [M-2H-H_3PO_3{}^{18}O]^{2-}$ | 10 | 3 |
| $[^{18}O_4]$ $InsP_7$ | 372.9 | 321.9 | $[M-2H]^{2-} \rightarrow [M-2H-H_3PO_2{}^{18}O_2]^{2-}$ | 10 | 3 |
| $[^{18}O_4]$ $InsP_7$ | 372.9 | 320.9 | $[M-2H]^{2-} \rightarrow [M-2H-H_3PO{}^{18}O_3]^{2-}$ | 10 | 3 |
| $[^{18}O_4]$ $InsP_7$ | 372.9 | 319.9 | $[M-2H]^{2-} \rightarrow [M-2H-H_3P{}^{18}O_4]^{2-}$ | 10 | 3 |
| $[^{18}O_5]$ $InsP_7$ | 373.9 | 324.9 | $[M-2H]^{2-} \rightarrow [M-2H-H_3PO_4]^{2-}$ | 10 | 3 |
| $[^{18}O_5]$ $InsP_7$ | 373.9 | 323.9 | $[M-2H]^{2-} \rightarrow [M-2H-H_3PO_3{}^{18}O]^{2-}$ | 10 | 3 |
| $[^{18}O_5]$ $InsP_7$ | 373.9 | 322.9 | $[M-2H]^{2-} \rightarrow [M-2H-H_3PO_2{}^{18}O_2]^{2-}$ | 10 | 3 |
| $[^{18}O_5]$ $InsP_7$ | 373.9 | 321.9 | $[M-2H]^{2-} \rightarrow [M-2H-H_3PO{}^{18}O_3]^{2-}$ | 10 | 3 |
| $[^{18}O_5]$ $InsP_7$ | 373.9 | 320.9 | $[M-2H]^{2-} \rightarrow [M-2H-H_3P{}^{18}O_4]^{2-}$ | 10 | 3 |
| $[^{18}O_6]$ $InsP_7$ | 374.9 | 325.9 | $[M-2H]^{2-} \rightarrow [M-2H-H_3PO_4]^{2-}$ | 10 | 3 |
| $[^{18}O_6]$ $InsP_7$ | 374.9 | 324.9 | $[M-2H]^{2-} \rightarrow [M-2H-H_3PO_3{}^{18}O]^{2-}$ | 10 | 3 |
| $[^{18}O_6]$ $InsP_7$ | 374.9 | 323.9 | $[M-2H]^{2-} \rightarrow [M-2H-H_3PO_2{}^{18}O_2]^{2-}$ | 10 | 3 |
| $[^{18}O_6]$ $InsP_7$ | 374.9 | 322.9 | $[M-2H]^{2-} \rightarrow [M-2H-H_3PO{}^{18}O_3]^{2-}$ | 10 | 3 |
| $[^{18}O_6]$ $InsP_7$ | 374.9 | 321.9 | $[M-2H]^{2-} \rightarrow [M-2H-H_3P{}^{18}O_4]^{2-}$ | 10 | 3 |

Notes: Negative polarity; fragmentor (V) is 166; dwell is 25.

Supplementary Table S8. MRM transitions setting of $InsP_6$ for amoeba samples

| Molecular name | Precursor Ion | Product Ion | Type of transition | Collision Energy (V) | Cell Accelerator Voltage |
|---|---|---|---|---|---|
| $InsP_6$ | 328.9 | 79 | $[M\text{-}2H]^{2-} \rightarrow [PO_3]^-$ | 13 | 4 |
| [$^{18}O$] $InsP_6$ | 329.9 | 81 | $[M\text{-}2H]^{2-} \rightarrow [PO_2{}^{18}O]^-$ | 13 | 4 |
| [$^{18}O$] $InsP_6$ | 329.9 | 79 | $[M\text{-}2H]^{2-} \rightarrow [PO_3]^-$ | 13 | 4 |
| [$^{18}O_2$] $InsP_6$ | 330.9 | 83 | $[M\text{-}2H]^{2-} \rightarrow [PO^{18}O_2]^-$ | 13 | 4 |
| [$^{18}O_2$] $InsP_6$ | 330.9 | 81 | $[M\text{-}2H]^{2-} \rightarrow [PO_2{}^{18}O]^-$ | 13 | 4 |
| [$^{18}O_2$] $InsP_6$ | 330.9 | 79 | $[M\text{-}2H]^{2-} \rightarrow [PO_3]^-$ | 13 | 4 |
| [$^{18}O_3$] $InsP_6$ | 331.9 | 85 | $[M\text{-}2H]^{2-} \rightarrow [P^{18}O_3]^-$ | 13 | 4 |
| [$^{18}O_3$] $InsP_6$ | 331.9 | 83 | $[M\text{-}2H]^{2-} \rightarrow [PO^{18}O_2]^-$ | 13 | 4 |
| [$^{18}O_3$] $InsP_6$ | 331.9 | 81 | $[M\text{-}2H]^{2-} \rightarrow [PO_2{}^{18}O]^-$ | 13 | 4 |
| [$^{18}O_3$] $InsP_6$ | 331.9 | 79 | $[M\text{-}2H]^{2-} \rightarrow [PO_3]^-$ | 13 | 4 |
| [$^{18}O_4$] $InsP_6$ | 332.9 | 85 | $[M\text{-}2H]^{2-} \rightarrow [P^{18}O_3]^-$ | 13 | 4 |
| [$^{18}O_4$] $InsP_6$ | 332.9 | 83 | $[M\text{-}2H]^{2-} \rightarrow [PO^{18}O_2]^-$ | 13 | 4 |
| [$^{18}O_4$] $InsP_6$ | 332.9 | 81 | $[M\text{-}2H]^{2-} \rightarrow [PO_2{}^{18}O]^-$ | 13 | 4 |
| [$^{18}O_4$] $InsP_6$ | 332.9 | 79 | $[M\text{-}2H]^{2-} \rightarrow [PO_3]^-$ | 13 | 4 |
| [$^{18}O_5$] $InsP_6$ | 333.9 | 85 | $[M\text{-}2H]^{2-} \rightarrow [P^{18}O_3]^-$ | 13 | 4 |
| [$^{18}O_5$] $InsP_6$ | 333.9 | 83 | $[M\text{-}2H]^{2-} \rightarrow [PO^{18}O_2]^-$ | 13 | 4 |
| [$^{18}O_5$] $InsP_6$ | 333.9 | 81 | $[M\text{-}2H]^{2-} \rightarrow [PO_2{}^{18}O]^-$ | 13 | 4 |
| [$^{18}O_5$] $InsP_6$ | 333.9 | 79 | $[M\text{-}2H]^{2-} \rightarrow [PO_3]^-$ | 13 | 4 |
| [$^{18}O_6$] $InsP_6$ | 334.9 | 85 | $[M\text{-}2H]^{2-} \rightarrow [P^{18}O_3]^-$ | 13 | 4 |
| [$^{18}O_6$] $InsP_6$ | 334.9 | 83 | $[M\text{-}2H]^{2-} \rightarrow [PO^{18}O_2]^-$ | 13 | 4 |
| [$^{18}O_6$] $InsP_6$ | 334.9 | 81 | $[M\text{-}2H]^{2-} \rightarrow [PO_2{}^{18}O]^-$ | 13 | 4 |
| [$^{18}O_6$] $InsP_6$ | 334.9 | 79 | $[M\text{-}2H]^{2-} \rightarrow [PO_3]^-$ | 13 | 4 |
| [$^{18}O_7$] $InsP_6$ | 335.9 | 85 | $[M\text{-}2H]^{2-} \rightarrow [P^{18}O_3]^-$ | 13 | 4 |
| [$^{18}O_7$] $InsP_6$ | 335.9 | 83 | $[M\text{-}2H]^{2-} \rightarrow [PO^{18}O_2]^-$ | 13 | 4 |
| [$^{18}O_7$] $InsP_6$ | 335.9 | 81 | $[M\text{-}2H]^{2-} \rightarrow [PO_2{}^{18}O]^-$ | 13 | 4 |
| [$^{18}O_7$] $InsP_6$ | 335.9 | 79 | $[M\text{-}2H]^{2-} \rightarrow [PO_3]^-$ | 13 | 4 |
| [$^{18}O_8$] $InsP_6$ | 336.9 | 85 | $[M\text{-}2H]^{2-} \rightarrow [P^{18}O_3]^-$ | 13 | 4 |
| [$^{18}O_8$] $InsP_6$ | 336.9 | 83 | $[M\text{-}2H]^{2-} \rightarrow [PO^{18}O_2]^-$ | 13 | 4 |
| [$^{18}O_8$] $InsP_6$ | 336.9 | 81 | $[M\text{-}2H]^{2-} \rightarrow [PO_2{}^{18}O]^-$ | 13 | 4 |
| [$^{18}O_8$] $InsP_6$ | 336.9 | 79 | $[M\text{-}2H]^{2-} \rightarrow [PO_3]^-$ | 13 | 4 |

Notes: Negative polarity; fragmentor (V) is 166; dwell is 30.

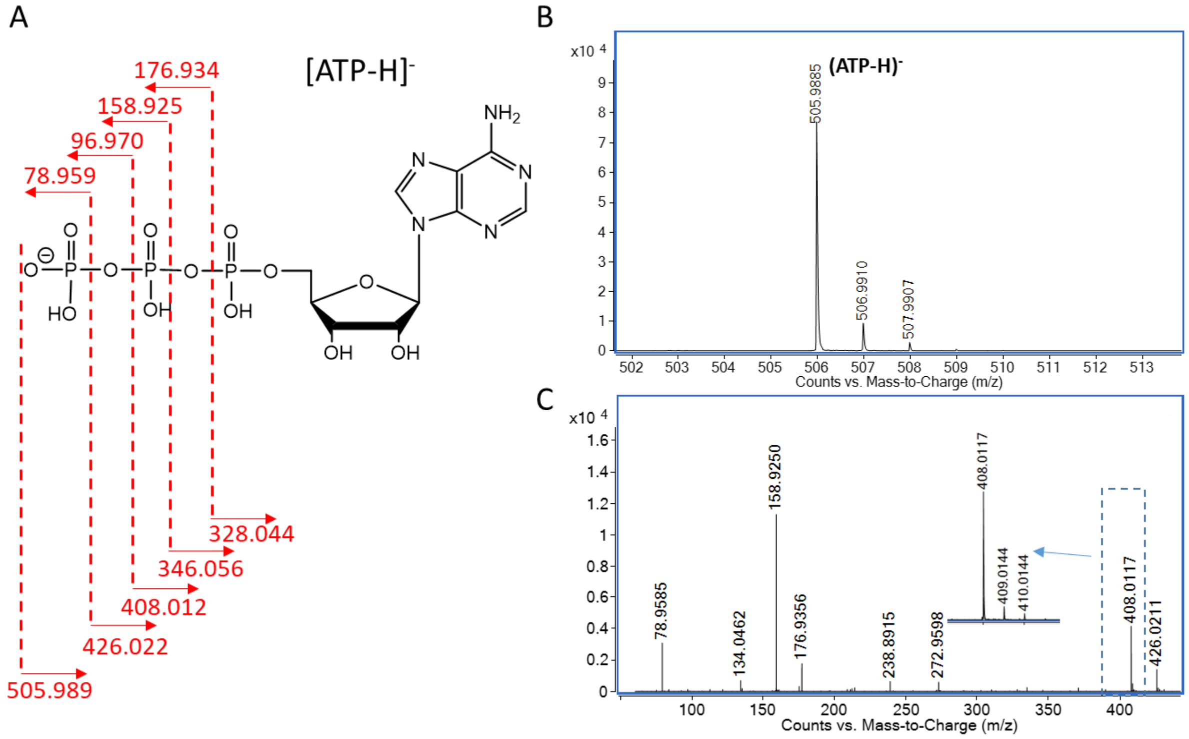


**Supplementary Figure S1.** ESI-MS analysis of ATP ESI-MS fragmentations of ATP and proposed assignment of the number of $^{18}O$ labels on the γ-phosphate. **A** Structure and proposed ESI-MS fragmentations of ATP. **B** ESI-MS of ATP standard. **C** Observed ESI-MS fragmentations of ATP by qTOF. 408.0117, which corresponds to (ATP-H-$H_3PO_4$)-, was taken as the special product ion for the QQQ analysis. The number of the $^{18}O$ labels on the γ-phosphate was assigned according to the $^{18}O$ labeling of this special product ion. For example, ($^{18}O_1$ ATP-H-$H_3PO_4$)-was assigned to be one $^{18}O$ labeling on γ-phosphate and ($^{18}O_1$ ATP-H-$H_3PO_3{}^{18}O$)$^-$ was assigned to be zero $^{18}O$ labeling on γ-phosphate.

A

B

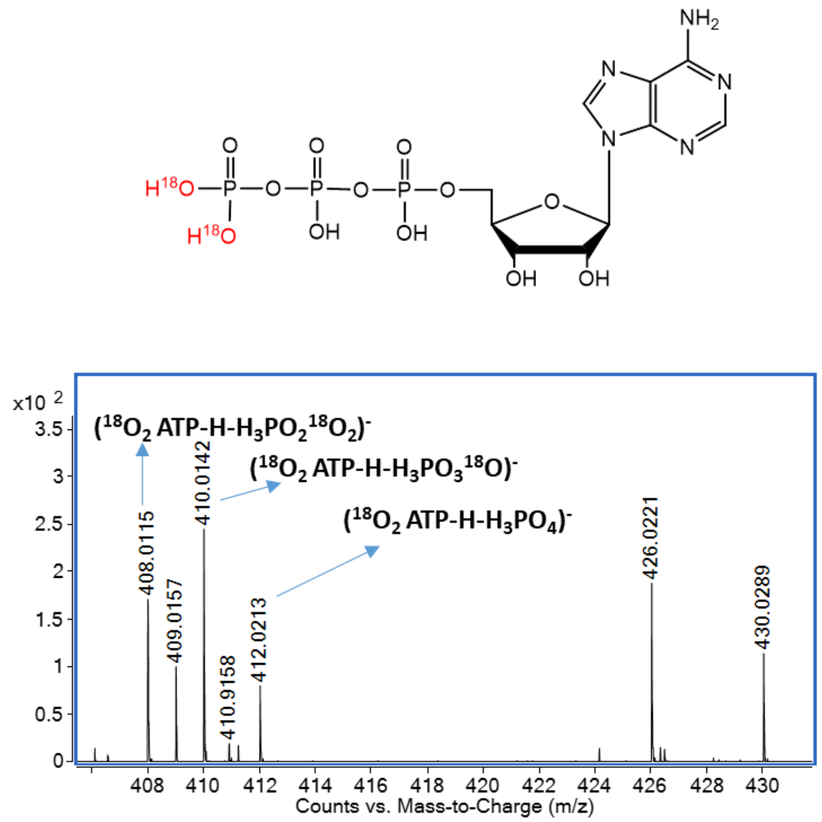


**Supplementary Figure S2.** Oxygen migration (scrambling) on ATP in the gas phase. **A** Structure of synthetic γ-$^{18}O_2$ labeled ATP. **B** ($^{18}O_2$ ATP-H-$H_3PO_2{}^{18}O_2$)$^-$, ($^{18}O_2$ ATP-H-$H_3PO_3{}^{18}O$)$^-$ and ($^{18}O_2$ ATP-H-$H_3PO_4$)$^-$ were observed, indicating partial scrambling of $^{18}O$ in the gas phase (qTOF).

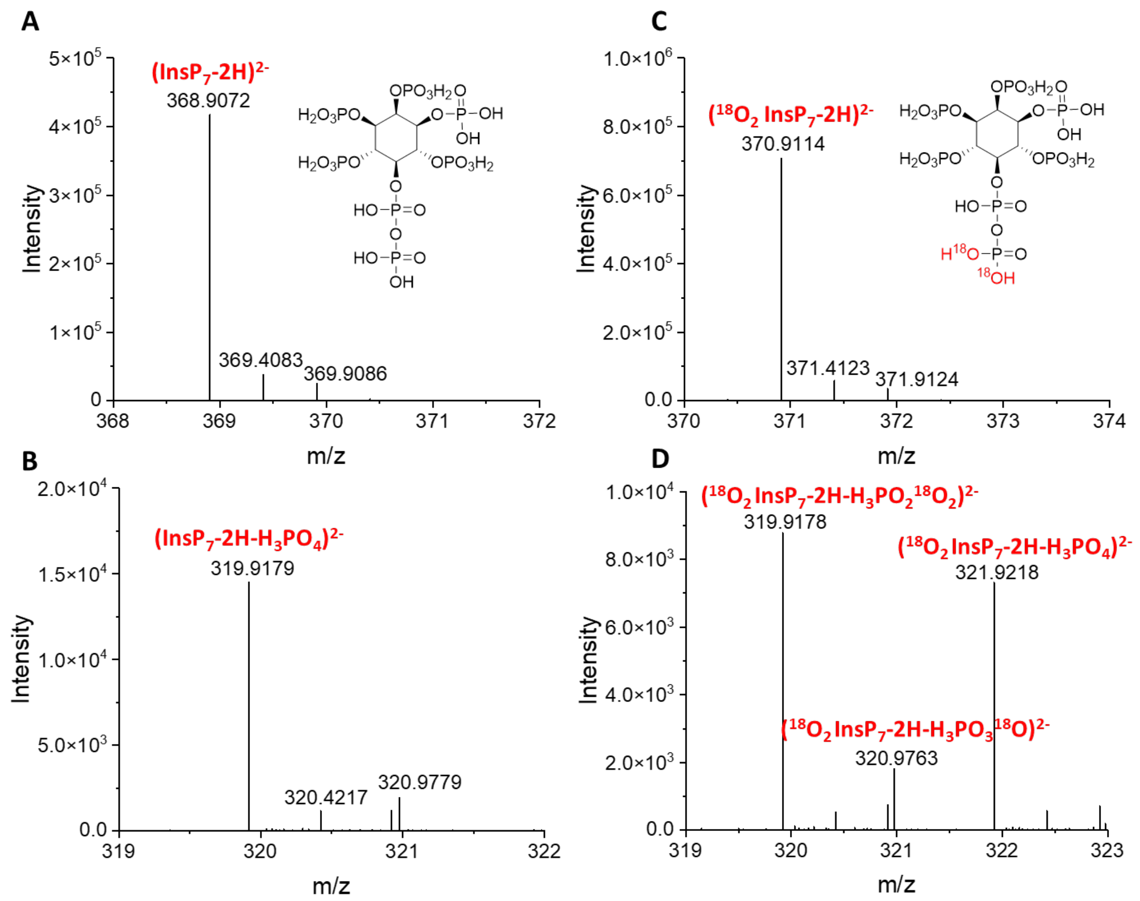


**Supplementary Figure S3.** Oxygen migration (scrambling) on 5-$InsP_7$ in the gas phase. **A** ESI-MS of 5-$InsP_7$ standard. **B** Observed ESI-MS fragmentations of 5-$InsP_7$ standard by qTOF. **C** ESI-MS of synthetic $^{18}O_2$ 5-$InsP_7$ standard. **D** Observed ESI-MS fragmentations of synthetic $^{18}O_2$ 5-$InsP_7$ standard by qTOF. The results show that partial scrambling of $^{18}O$ in the gas phase is also occurring in $InsP_7$.

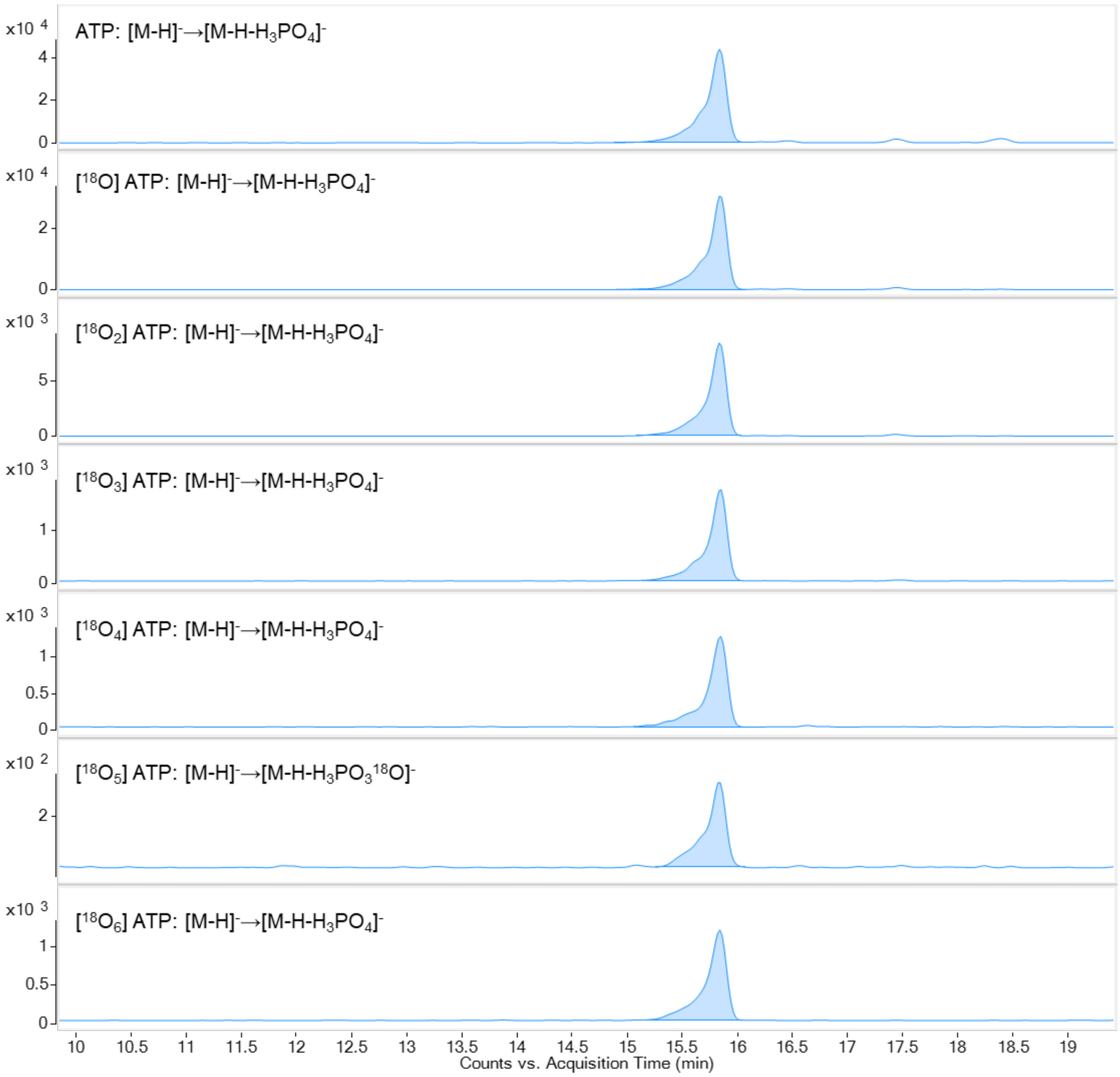


**Supplementary Figure S4.** Extracted ion electropherograms (EIEs) of unlabeled and $^{18}O$ labeled ATP from yeast under steady state conditions at the 1 min time point.

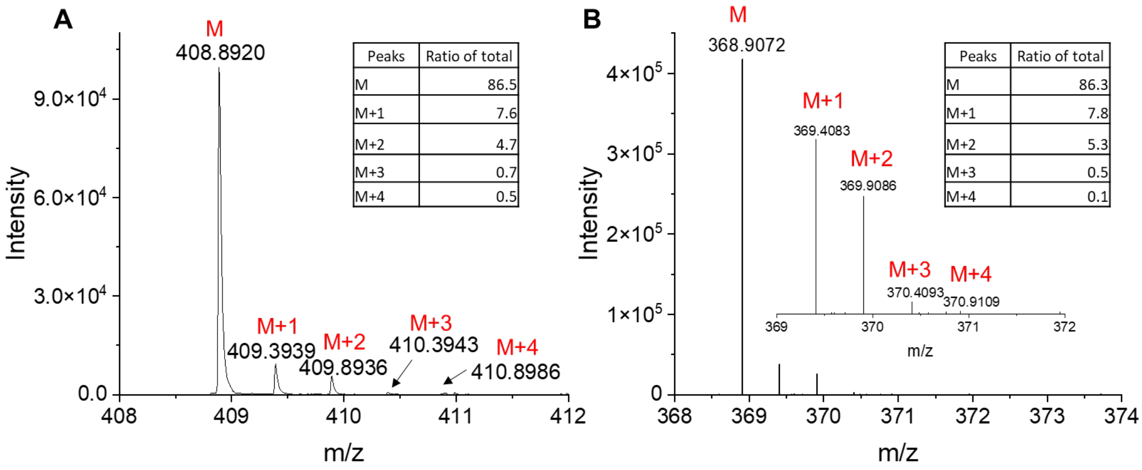


**C**

| | Wide mass resolution (±0.6 Da mass error) | | | Unit mass resolution (±0.35 Da mass error) | | |
|---|---|---|---|---|---|---|
| | % of total | | | % of total | | |
| | $^{18}O_0$ (M) | $^{18}O_1$ (M+2) | $^{18}O_2$ (M+4) | $^{18}O_0$ (M) | $^{18}O_1$ (M+2) | $^{18}O_2$ (M+4) |
| 1,5-Ins$P_8$ | 87.9 | 11.6 | 0.5 | 94.0 | 5.8 | 0.2 |
| 5-Ins$P_7$ | 88.4 | 11.2 | 0.4 | 94.4 | 5.6 | 0.04 |

**Supplementary Figure S5.** Effect of the isotopic peak. **A** Distribution of isotopic species in unlabeled 1,5-Ins$P_8$ was detected by qTOF in the state of double charged ion. **B** Distribution of isotopic species in unlabeled 5-Ins$P_7$ was detected by qTOF in the status of double charged ion. Thus, the delta mass between M and M+1 is 0.5 Da. **C** Unlabeled 1,5-Ins$P_8$ and 5-Ins$P_7$ were detected by CE-QQQ system with both wide mass resolution and unit mass resolution. Unit mass resolution is able to resolve M, M+1, M+2, M+3, and M+4, because the unit mass error is ± 0.35 Da, which is narrower than the delta mass of doubly charged isotopic species (0.5 Da). Nevertheless M+1 and M+2 cannot be resolved by wide mass resolution (0.6 Da mass error). Consequently, for wide resolution, detected $^{18}O_1$ (M+2) contains M+1, M+2 or even M+3, corresponding to ca. 12% of total 1,5-Ins$P_8$ or ca. 11% of total 5-Ins$P_7$. The ratio of isotopic species found by qTOF is matched by this data.

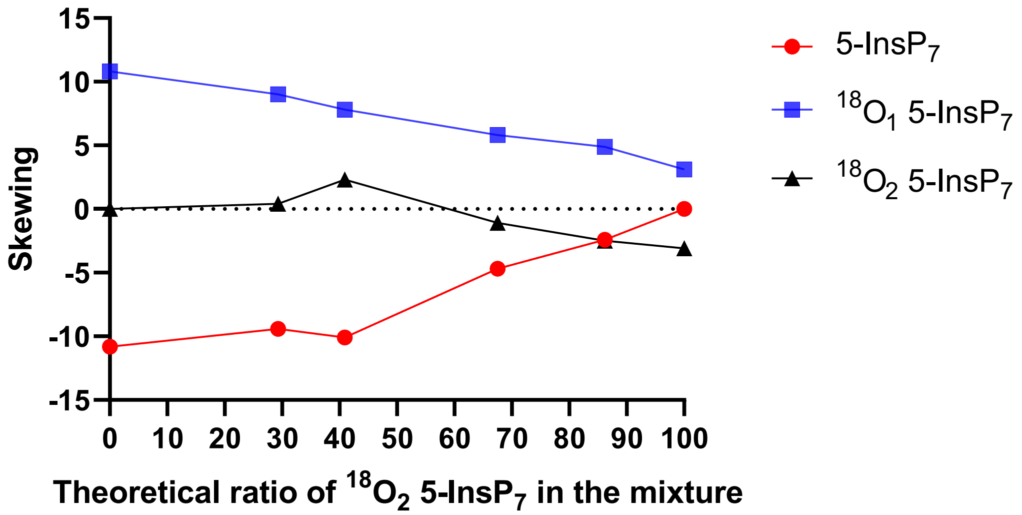


**Supplementary Figure S6.** Mixtures of synthetic $^{18}O_2$ 5-InsP$_7$ and unlabeled 5-InsP$_7$ with different compositions according to the theoretical ratio were prepared and analyzed by CE-QQQ. Skewing = experimental ratio (by CE-QQQ) - theoretical ratio.

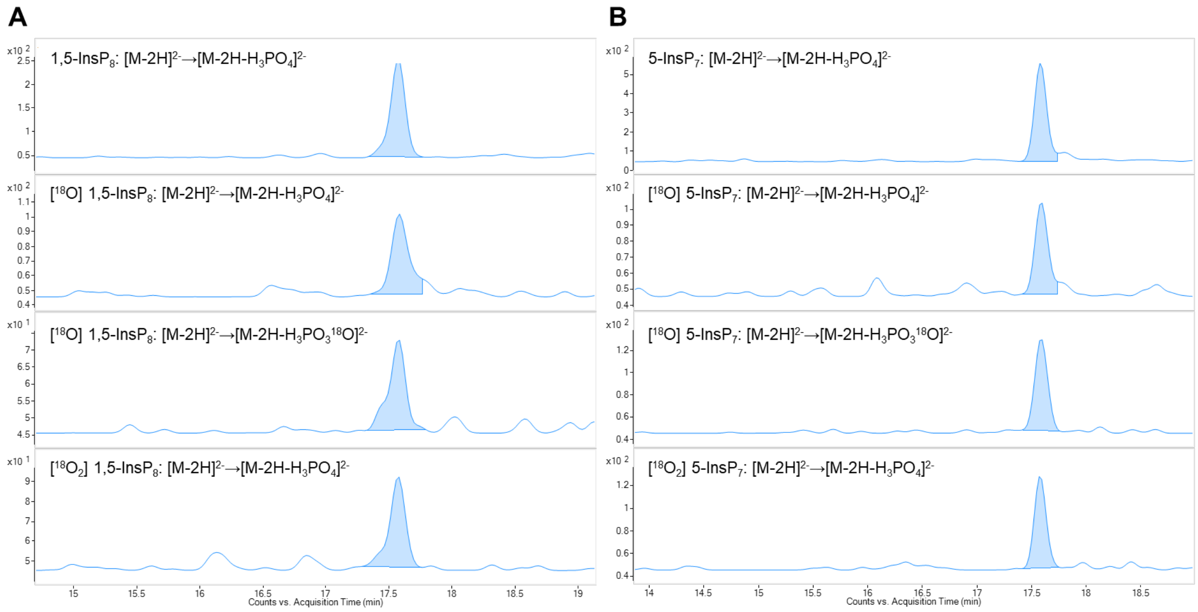


**Supplementary Figure S7.** Extracted ion electropherograms (EIEs) of unlabeled and $^{18}$O labeled 1,5-InsP$_8$ (**A**) and 5-InsP$_7$ (**B**) from yeast under steady state conditions at the 1 min time point.

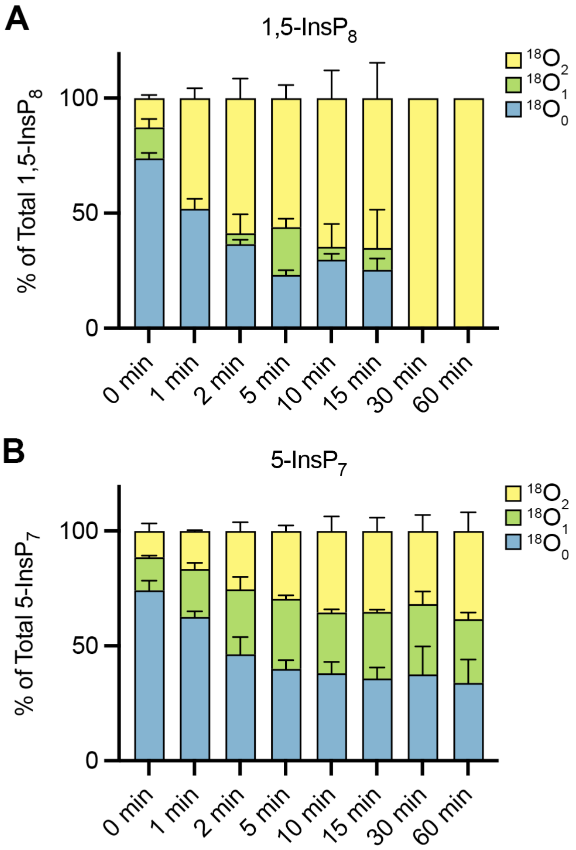


**Supplementary Figure S8.** Kinetics of $^{18}O$ entry into soluble InsPs of yeast under steady state conditions. Wild type cells were grown logarithmically in SC medium. The medium was changed to SC medium prepared with 100% of $^{18}O$-labelled water. After further incubation at 20˚C for the indicated time periods, cells were extracted with perchloric acid. The means of triplicates are shown with standard deviation, representing: **A**. 1,5-InsP$_8$, **B**. 5-InsP$_7$.

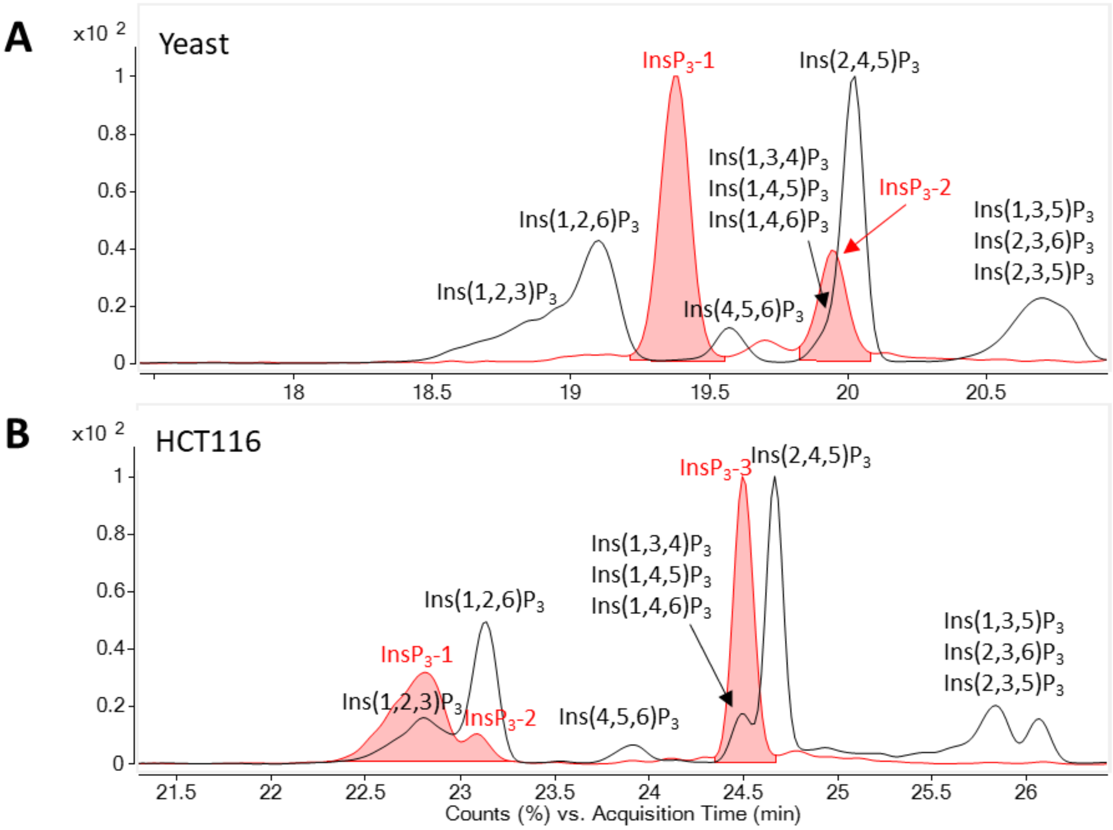


**Supplementary Figure S9** Assignment of $InsP_3$ isomers in yeast and HCT116 cells. **A** Extracted ion electropherograms of [$^{13}C_6$] $InsP_3$ reference (black line) and $InsP_3$-1 and $InsP_3$-2 in yeast (red area). The [$^{13}C_6$] $InsP_3$ reference were generated by incubating [$^{13}C_6$] $InsP_6$ (prepared with ultrapure water) at 100°C for 5 h. The assignment of each $InsP_3$ isomers were achieved as previously described (Liu *et al*, 2023). $InsP_3$-2 comigrate with [$^{13}C_6$] Ins(1,3,4)$P_3$, Ins(1,4,5)$P_3$, Ins(1,4,6)$P_3$ and/or its enantiomers, and we infer that $InsP_3$-2 is Ins(1,4,5)$P_3$. **B** Extracted ion electropherograms of [$^{13}C_6$] $InsP_3$ reference (black line) and $InsP_3$-1, $InsP_3$-2 and $InsP_3$-3 in HCT116 cells (red area). $InsP_3$-1 comigrates with [$^{13}C_6$] Ins(1,2,3)$P_3$ and $InsP_3$-2 comigrates with [$^{13}C_6$] Ins(1,2,6)$P_3$ and/or its enantiomers. $InsP_3$-3 comigrates with [$^{13}C_6$] Ins(1,3,4)$P_3$, Ins(1,4,5)$P_3$, Ins(1,4,6)$P_3$ and/or its enantiomers, and from we infer that $InsP_3$-3 is Ins(1,4,5)$P_3$ as most likely species.

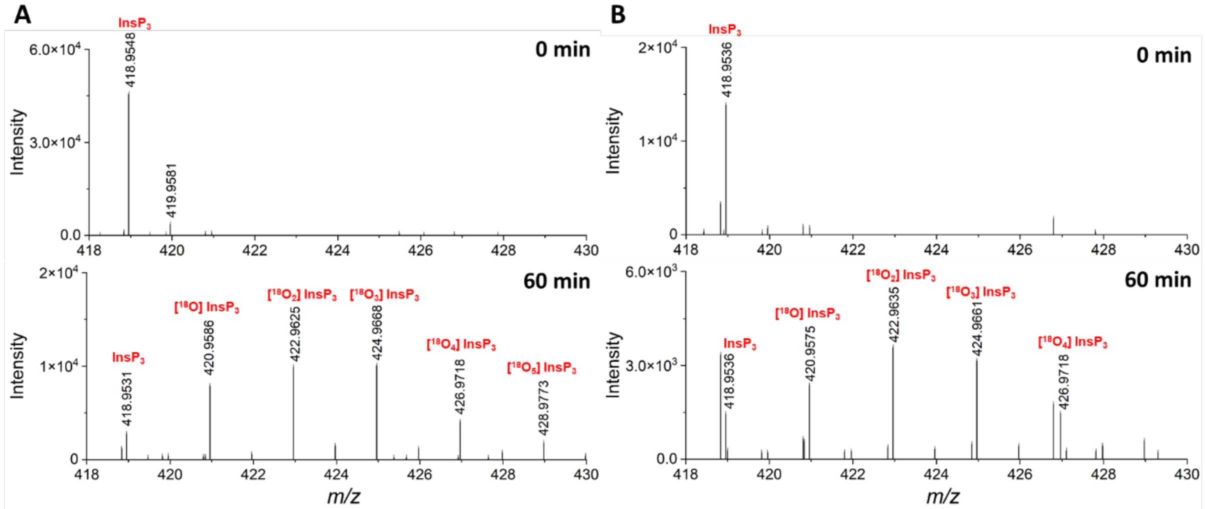


**Supplementary Figure S10** The qTOF analysis of $InsP_3$ in yeast. The analysis reveals the kinetics of $^{18}O$ incorporation into $InsP_3$-1 (**A**) and $InsP_3$-2 (**B**) at 0 min and 60 min time point. Theoretical $[M\text{-}H]^-$ for $InsP_3$, [$^{18}O$] $InsP_3$, [$^{18}O_2$] $InsP_3$, [$^{18}O_3$] $InsP_3$, [$^{18}O_4$] $InsP_3$, and [$^{18}O_5$] $InsP_3$ is 418.9551, 420.9593, 422.9636, 424.9678, 426.9721, 428.9763, respectively.

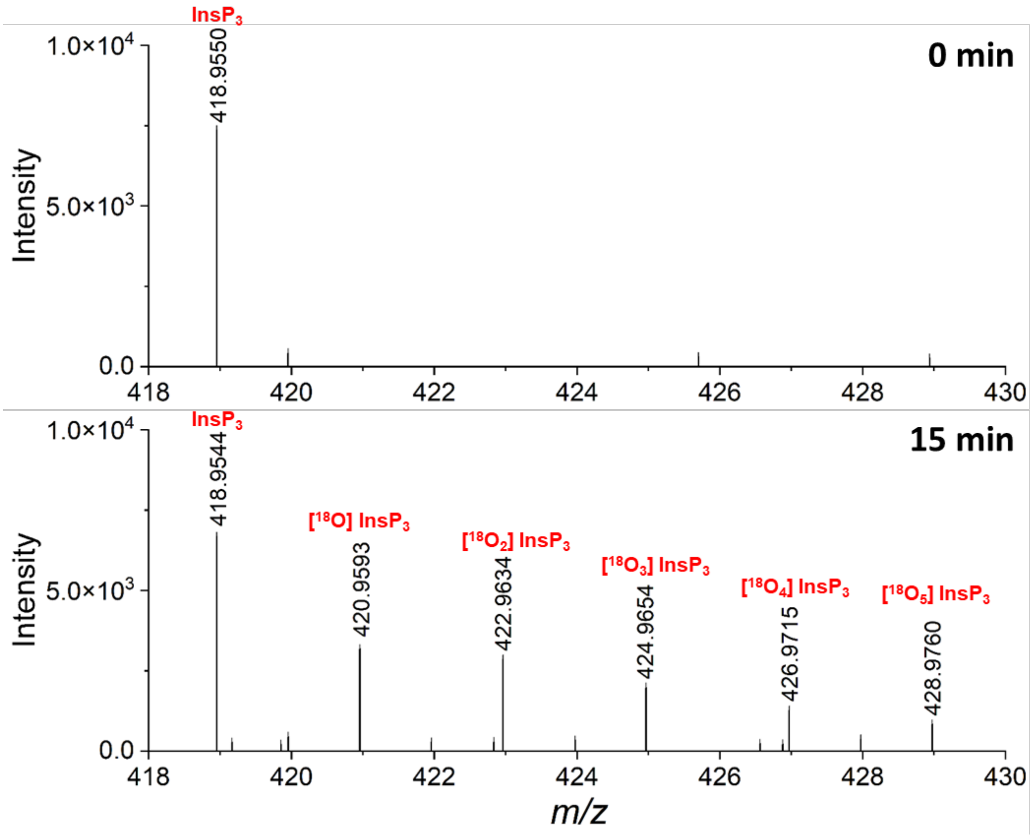


**Supplementary Figure S11** The qTOF analysis of $InsP_3$-3 in HCT116 cells. The analysis reveals the kinetics of $^{18}O$ incorporation into $InsP_3$-3 at 0 min and 15 min time point. Theoretical $[M-H]^-$ for $InsP_3$, [$^{18}O$] $InsP_3$, [$^{18}O_2$] $InsP_3$, [$^{18}O_3$] $InsP_3$, [$^{18}O_4$] $InsP_3$, and [$^{18}O_5$] $InsP_3$ is 418.9551, 420.9593, 422.9636, 424.9678, 426.9721, 428.9763, respectively.

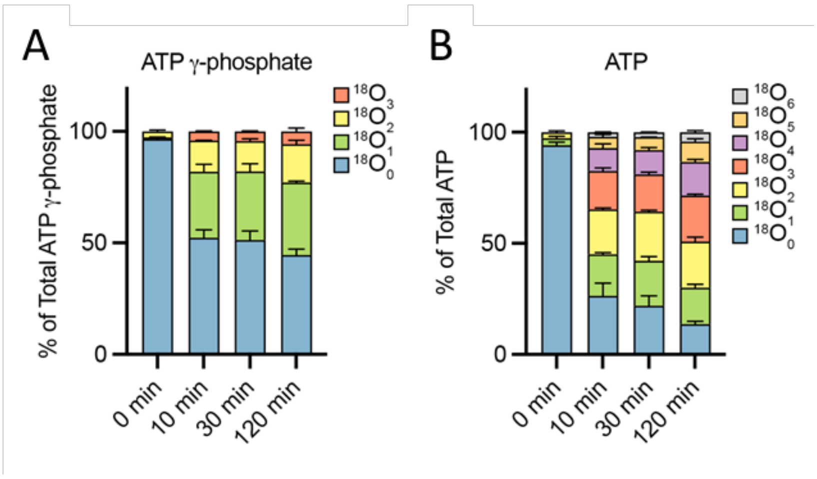


**Supplementary Figure S12.** Kinetics of $^{18}$O entry into ATP from *D. discoideum*. Kinetics of $^{18}$O entry into ATP γ-phosphate (**A**) and ATP (**B**) were monitored in *D. discoideum*. Cells were grown in SIH medium, transferred to SIH media made of 50% of $^{18}$O-labeled water. After further incubation for the indicated periods of time, samples were harvested and extracted. The means of two replicates with deviations are shown.
